# Reported transgenerational responses to *Pseudomonas aeruginosa* in *C. elegans* are not robust

**DOI:** 10.1101/2024.06.01.596941

**Authors:** D. Patrick Gainey, Andrey V. Shubin, Craig P. Hunter

## Abstract

Here we report our attempt to replicate reports of transgenerational epigenetic inheritance in *Caenorhabditis elegans*. Published results from multiple laboratories show that *C. elegans* adults and their F1 embryos exposed to the pathogen *Pseudomonas aeruginosa* show pathogen aversion behavior and a pathogen exposure-dependent increase in *daf-7/TGFβ* reporter gene expression. However, results from one group show persistence of the aversion behavior and elevated *daf-7* expression in the F2, F3, and F4 generations. In contrast, we failed to consistently detect either the pathogen avoidance response or elevated *daf-7* expression beyond the F1 generation. We did confirm that the dsRNA transport proteins SID-1 and SID-2 are required for the intergenerational (F1) inheritance of pathogen avoidance, but not for the F1 inheritance of elevated *daf-7* expression. Furthermore, our reanalysis of RNA seq data provides additional evidence that this intergenerational inherited PA14 response may be mediated by small RNAs. The experimental methods are well-described, the source materials are readily available, including samples from the reporting laboratory, and we explored a variety of environmental conditions likely to account for lab-to-lab variability. None of these adjustments altered our results. We conclude that this example of transgenerational inheritance lacks robustness, confirm that the intergenerational avoidance response, but not the elevated *daf-7p::gfp* expression in F1 progeny, requires *sid-1* and *sid-2*, and identify candidate siRNAs and target genes that may mediate this intergenerational response.

## Introduction

*C. elegans* worms exposed to the pathogen *Pseudomonas aeruginosa* (strain PA14) learn to avoid this specific pathogen upon subsequent exposure (Zhang et al., 2005). PA14 exposure also induces the expression of a reporter (*daf-7p::gfp*) for the cytokine TGF-β in the ASI neurons of PA14-exposed animals (Meisel and Kim, 2014). In Moore et al., (2019) and three follow-up papers (Kaletsky et al., 2020; Moore et al., 2021a; Sengupta et al., 2024), Murphy and colleagues report that *C. elegans* adults trained to avoid PA14 transmit this learned avoidance and elevated *daf-7p::gfp* expression in ASI neurons to four generations of progeny (F1-F4). They further reported that numerous RNA interference (RNAi) factors, including the dsRNA transport proteins SID-1 and SID-2 (Winston et al., 2002; Feinberg 2003; Winston et al., 2007; McEwan et al., 2012), are required for the inheritance of the behavioral response and elevated *daf-7p::gfp* expression (Kaletsky et al., 2020; Moore et al., 2019). To date, no follow-up reports by independent groups have been published on the transgenerational character of this response. We were keen to reproduce these results and investigate in detail the contributions of the dsRNA transport proteins SID-1 and SID-2 to transgenerational epigenetic inheritance (TEI).

We readily reproduced learned behavior and elevated *daf-7p::gfp* expression in trained parents (P0) and their F1 progeny, but after many attempts and numerous protocol adjustments, we failed to reproducibly replicate inheritance among F2 progeny. While we have been unable to identify a specific methodological cause for our different results, we conclude that this example of TEI is insufficiently robust for experimental investigation.

## Results and Discussion

### The reported PA14 training conditions failed to produce an avoidance response or elevated *daf-7p::gfp* expression in ASI neurons among F2 progeny

PA14-exposed animals (P0) and their progeny (F1) avoid PA14 in subsequent choice assays and show elevated *daf-7p::gfp* expression in ASI neurons (Zhang et al., 2005; Moore et al., 2019; Kaletsky et al., 2020; Pereira et al., 2020; Sengupta et al., 2024). The Murphy group has reported that both responses are transgenerationally inherited by F2-F4 generation worms (Moore et al., 2019; Kaletsky et al., 2020; Moore et al., 2021a; Sengupta et al., 2024). Assays for transgenerational inheritance in *C. elegans*, like the choice assay described in Moore et al., (2019), are frequently based on the collective behavior of a population. In contrast, the heritable increase in *daf-7p::gfp* expression in ASI neurons is a single animal assay that can be reliably scored in a few dozen animals (Moore et al., 2019). Recognizing the potential of this single-animal assay to aid investigation of the reported RNAi-pathway dependent response to PA14 exposure, we initially attempted to replicate the reported transgenerational *daf-7p::gfp* expression results in the wild-type reporter strain. When these experiments failed to produce an F2 response, we then included the population-based choice assays to assist in troubleshooting.

We performed the choice assay and *daf-7p::gfp* expression assay experiments as described in the published protocols (Moore et al., 2019; Kaletsky et al., 2020; Moore et al., 2021b) with minor adjustments (see Methods and S1_aversion_protocol). The pathogen avoidance response in the trained animals (P0 generation) was robust and often detected among their F1 progeny fertilized during parental exposure (Figures 1A-1C). However, the transgenerational (F2) response was not detected (Figures 1A-1C). In contrast, while the magnitude and significance of the elevated *daf-7p::gfp* expression in the ASI neurons in the P0 generation was variable, the response in the F1 generation was strong and statistically highly significant (Figures 1D-1I). Unlike the results reported in Moore et al., (2019) and Kaletsky et al., (2020), we did not observe a *daf-7p::gfp* response in the F2 generation (Figures 1D-1I).

**Figure 1.**
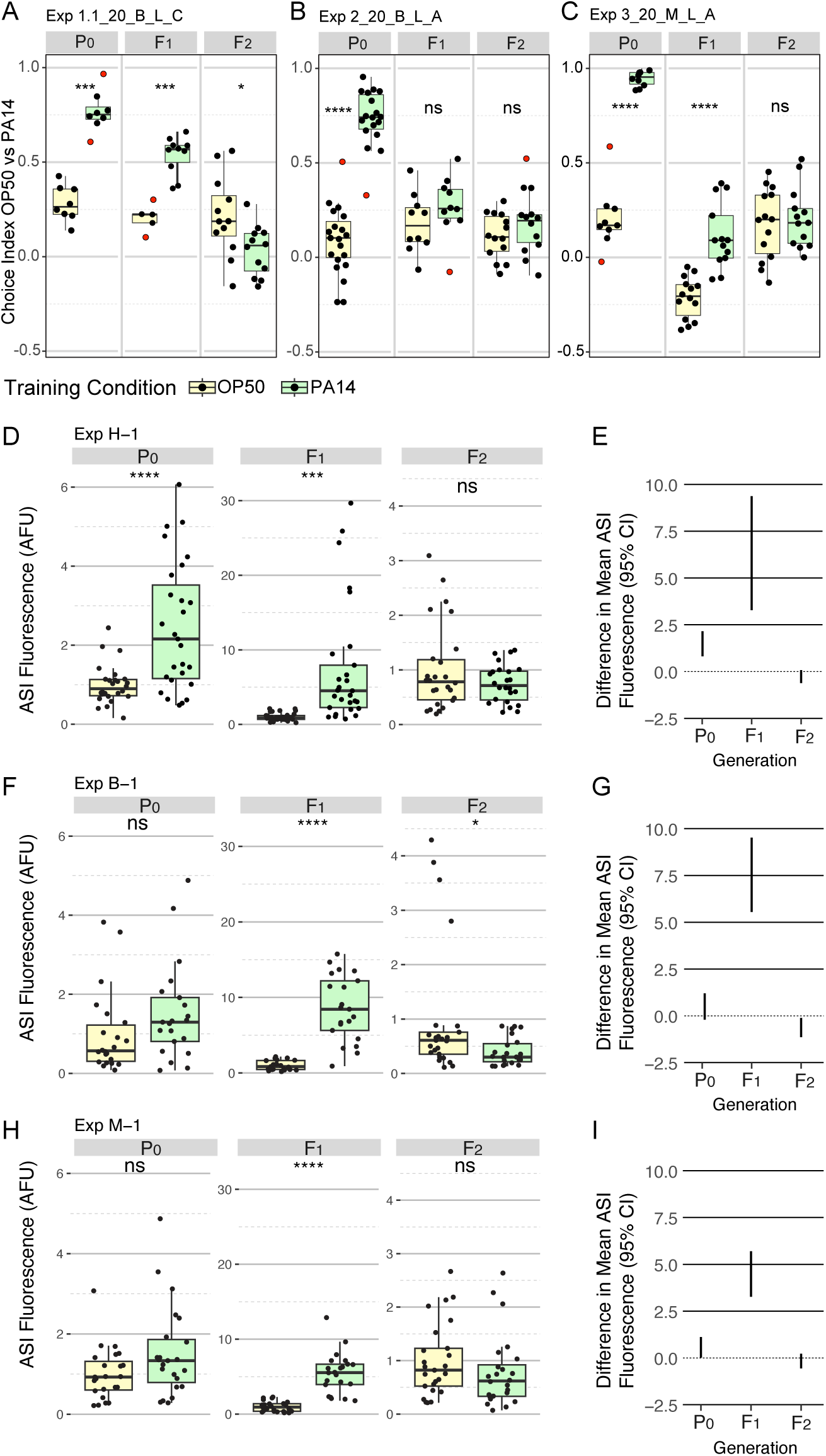
P0, F1 and F2 generation responses to P0 PA14 exposure. Three representative experimental results for trained and inherited aversion behavior (A-C) and induced and inherited elevated *daf-7p::gfp* in ASI neurons (D-I). A-C, Quartile box plots for each generation and training condition showing the distribution of choice index values ([number of animals choosing OP50 - number of animals choosing PA14] / total number of choices) for each OP50 vs PA14 choice assay plate following the published protocols. The experimental and assay conditions are indicated in the panel titles; [Exp #] [growth temperature (20°C or 25°C)] [PA14 isolate (B, Balskus or M, Murphy)] [light/dark assay condition (L or D)] [azide/cold paralytic (A or C)]. D, F, H, Quartile box plots displaying average ASI neuron *daf-7p::gfp* expression levels per worm and E, G, I, show the 95% confidence intervals for the difference in absolute mean between the conditions. For these experiments the FK181 strain (integrated multi-copy [MC] *daf-7p::gfp* reporter) was cultured at 20°C at all generations and was exposed to one of three different PA14 isolates (H, Hunter; B, Balskus; M, Murphy labs). AFU arbitrary fluorescence units normalized to mean OP50 levels. Red dots in choice assay results indicate outlier data points that were included in all statistical tests. Statical significance **** P < 0.0001, *** < 0.001, ** < 0.01, * < 0.05, ns > 0.05. See Methods section for statistical methods.

Motivated to reproduce the reported results, we obtained and tested independent isolates of the bacterial strains PA14 and OP50. We obtained and used a PA14 isolate from the Balskus lab (Harvard University) and both PA14 and OP50 isolates from the Murphy lab (Princeton University). Similar results were obtained using these reagents (Figure 1, Table 1, and Table 2).

**Table 1.** Experimental conditions and results for *daf-7p::gfp* expression levels in ASI neurons of wild-type animals.

| Reporter* | Pre-growth temperature | P0-F2 growth temperature | PA14 isolate† | Numerical index | F1 elevated *daf-7p::gfp* | F2 elevated *daf-7p::gfp* |
| --- | --- | --- | --- | --- | --- | --- |
| MC | 15°C | 20°C | B | 1, 2, 3, 4, 5 | Y | N |
| MC | 20°C | 20°C | B | 1, 2, 3 | Y | N |
| MC | 20°C | 20°C | H | 1** | Y | N |
| MC | 20°C | 20°C | M | 1, 2, 3 | Y | N |
| SC | 20°C | 20°C | B | 1 | Y | N |
| SC | 20°C | 20°C | H | 1 | Y | N |
| MC | 20°C | 25°C | B | 1, 3 | Y | Y |
| MC | 20°C | 25°C | B | 2 | Y | N |
| SC | 20°C | 25°C | B | 1, 3 | Y | Y |
| SC | 20°C | 25°C | B | 2, 4 | Y | N |

\* MC, multi-copy reporter in strain FK181; SC, single-copy reporter in strain QL296
† B, Balskus lab isolate; H, Hunter lab isolate; M, Murphy lab isolate.
\*\* Prepared training plates were stored at 4°C for 48hrs prior to start of training

**Table 2.** Experimental conditions and results for PA14 avoidance (Choice) assay.

| Experiment | Genotype | Growth temp | PA14 isolate* | Light (L)<br>Dark (D) | Azide (A)<br>Cold (C) | F1 PA14 avoidance | F2 PA14 avoidance | Notes |
| --- | --- | --- | --- | --- | --- | --- | --- | --- |
| 1.1 | N2 | 20°C | B | L | C | Y | N | † |
| 1.2 | N2 | 20°C | B | D | C | NA | N | † |
| 2 | N2 | 20°C | B | L | A | N | N | # |
| 3 | N2 | 20°C | M | L | A | Y | N |  |
| 4.1 | N2 | 20°C | B | L | A | N | N | ** |
| 4.2 | N2 | 20°C | B | L | A | N | N | ** |
| 5.1 | N2 | 25°C | B | D | C | Y | N | †† |
| 5.2 | N2 | 25°C | B | D | C | N | Y | †† |
| 6 | N2 | 20°C | B | D | C | N | N |  |
| 7 | *sid-1(qt9)* | 20°C | B | D | C | N | NA |  |
| 8 | *sid-1(qt9)* | 25°C | B | L | C | N | NA |  |
| 9 | *sid-2(qt42)* | 20°C | B | D | C | N | NA |  |
| 10 | *sid-2(qt42)* | 25°C | B | L | C | N | NA |  |

\* B, Balskus lab isolate; M, Murphy lab isolate
† A single biological replicate was split and separately assayed in the light and dark

### Modifying training and growth conditions did not result in more reliable detection of transgenerational responses to PA14 exposure

To attempt to replicate these key observations, we then explored procedural and environmental variations. To train animals to recognize and avoid PA14, gravid wild-type (N2) adults grown on high growth (HG) plates seeded with *E. coli* strain OP50 were treated with sodium hypochlorite (bleach) to prepare aseptic P0 embryos, which were hatched and grown on HG OP50 plates at 20°C for 48-52 hours. Larval stage 4 (L4) animals were then gently washed from these plates and placed on normal growth (NG) plates seeded with either OP50 (control) or PA14 (training) for 24 hours and then similarly recovered for testing and bleach isolation of F1 embryos.

Because the 24-hour training period on PA14 limits reproduction and therefore recovery of F1 progeny, the published protocols (Moore et al., 2019; Moore et al., 2021b) suggest plating four times more animals on PA14 than on OP50 to compensate for the reduced fertility. We found this insufficient to reliably produce enough embryos for multigenerational experiments, and therefore also delayed the start of training until most animals were young adults (56-60 hours). The updated protocol for PA14 training also introduces a similar delay (52-56 hours post bleach) prior to training (C. Murphy personal communication). Importantly, training either late L4 animals (51-52 hours) or young adult animals (57-58 hours) both produced strong PA14 aversion responses in the trained populations (Figure 1 and Figure S1). Furthermore, a few *daf-7p::gfp* expression experiments that started with training P0 animals 48-54 hours post bleach were successfully completed. The results of these experiments did not differ from those obtained when training began with older animals (Figure 2, Table 1, Table S1). Training young adults greatly improved F1 egg recovery, allowing for more reliable testing of F1 and F2 progeny. Importantly, at the end of the 24-hour training period (80-84 hours since embryo isolation, 20°C) the naïve and trained worm populations had laid many eggs, ensuring that the recovered *in utero* F1 eggs were fertilized after exposure to PA14. This change in training time usually produced sufficient F1 and F2 animals for measurement of *daf-7p::gfp* expression levels in the ASI neurons and learned avoidance assays. However, the trained and control F2 progeny remained indistinguishable (Figure 1, Figure 2, Figure 3, Table 1, and Table 2).

**Figure 2.**
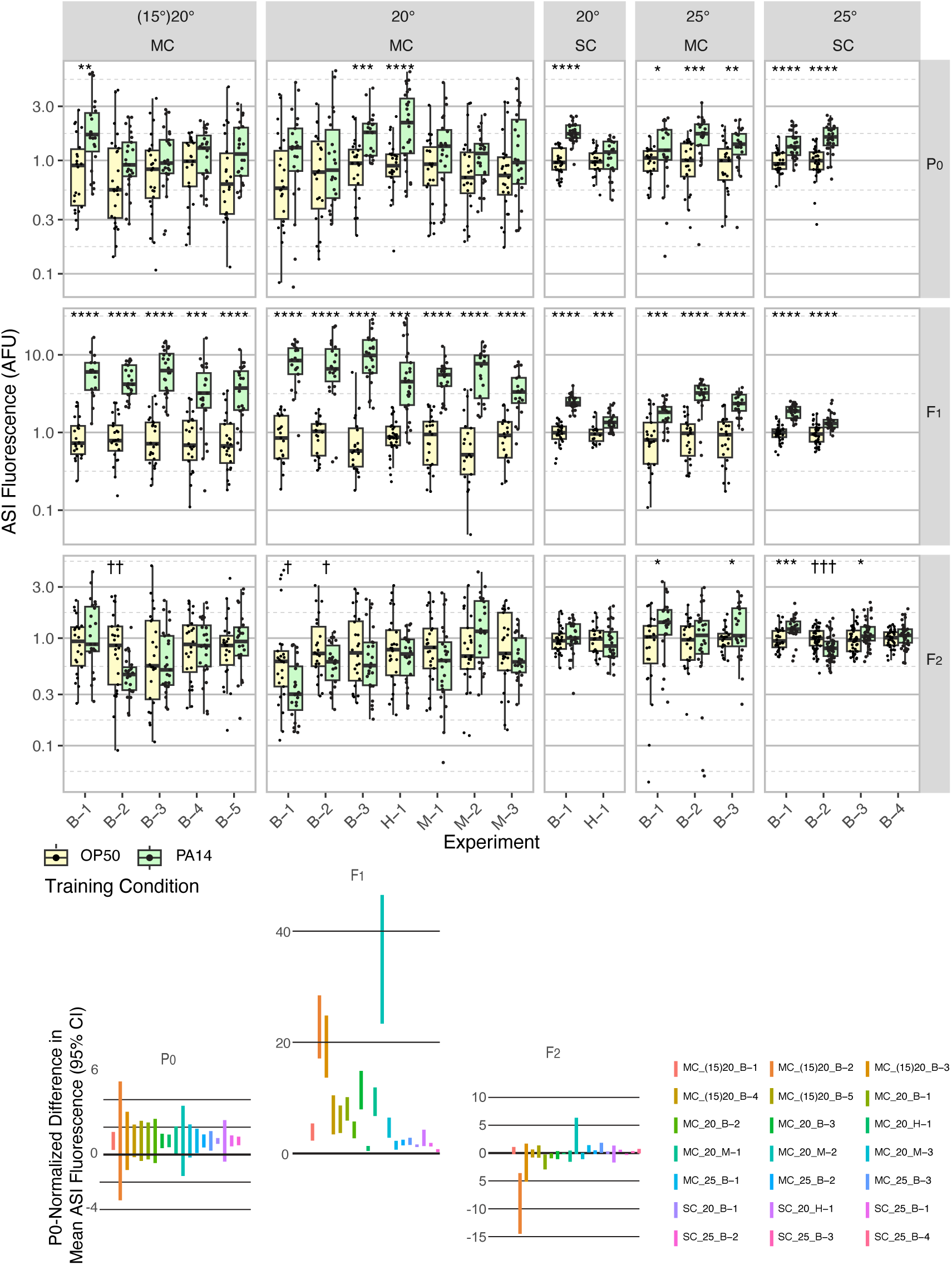
Effect of experimental design changes on multigenerational ASI *daf-7p::gfp* expression levels after P0 PA14 exposure. Box plots of results for 21 independent experiments are normalized to the average OP50 value by generation within each experiment for summary presentation (see Table 1 and Table S1 for experimental conditions). This figure includes the experiments shown in Figure 1. FK181 contains an integrated multi-copy (MC) tandem array composed of the *daf-7p::gfp* reporter and the co-injection marker *rol-6(su1006)* (Murakami et al., 2001). QL296 is a single copy (SC) insert of *daf-7p::gfp* with *unc-119(+)* as the co-selection marker (Zhan et al., 2015). Worms were cultured at either 20°C or 25°C and exposed to one of three different PA14 isolates (B, Balskus; H, Hunter; M, Murphy labs). In some experiments, worms were grown at 15°C for at least three generations prior to the P0 generation (indicated with parentheses, i.e. (15)20). The 95% confidence interval for the predicted absolute difference in means between conditions, normalized to the predicted P0 difference for each experiment (when applicable), is presented in the lower portion of the figure. Figure S2 shows the same data for individual neurons, rather than the mean of the two neurons. Statistical significance **** P < 0.0001, *** < 0.001, ** < 0.01, * < 0.05. Non-significant labels (p > 0.05) are omitted for clarity. † Indicate statistical significance with control higher than experiment. See Methods section for statistical methods.

**Figure 3.**
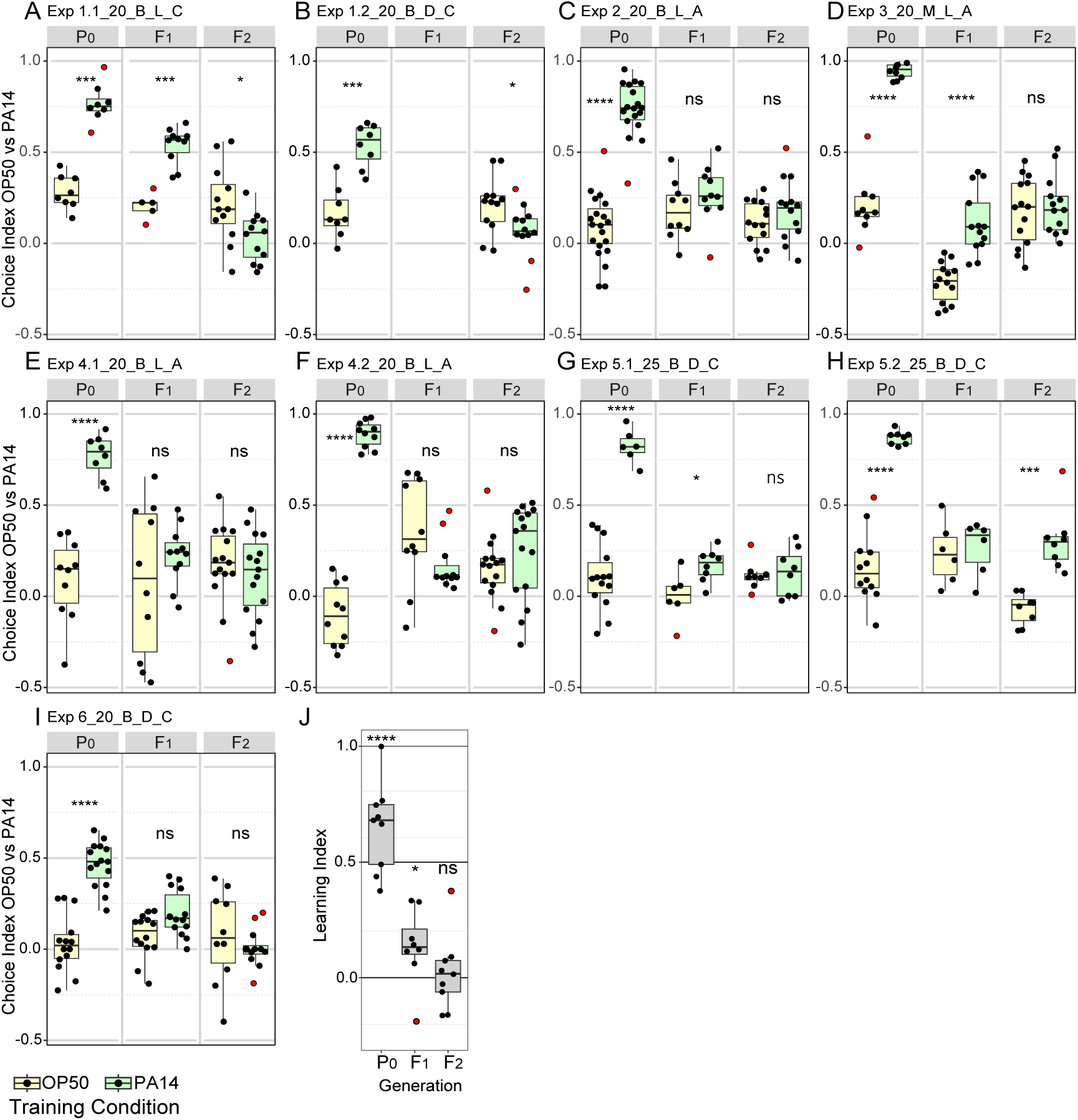
Effects of experimental design changes on multigenerational PA14 avoidance after PA14 exposure. Growth and assay conditions for each experiment are described in Table 2. This figure includes the experiments shown in Figure 1. Choice index calculated as described in Figure 1. A-I, The experimental and assay conditions are indicated in the panel titles; [Exp #] [growth temperature (20°C or 25°C)] [PA14 isolate (B, Balskus or M, Murphy)] [light/dark assay condition (L or D)] [azide/cold paralytic (A or C)]. J, Summary panel showing learning indexes (choice index for trained minus choice index for control) for all nine results (eight for F1 animals). See Figure S3 for Fisher’s exact test analysis of choice assay results. Red dots in choice assay results indicate outlier data points that were included in all statistical tests. Statical significance **** P < 0.0001, *** < 0.001,** < 0.01, * < 0.05, ns > 0.05. See Methods section for statistical methods.

To control for possible laboratory environmental differences, we also tested variations to the protocol, including worm growth temperature (15°C, 20°C, 25°C) before training (husbandry), during training, and in the F1-F2 generations (Figure 2, Figure 3, Table S1, and Table S2). These growth temperature variations were inspired by variability in reported husbandry conditions (Moore et al., 2019; Kaletsky et al., 2020; Moore et al., 2021b), the report that increased cultivation temperature (25°C) is known to increase *P. aeruginosa* pathogenicity (Tan et al., 1999), and the fact that the experiments that identified the *daf-7p::gfp* response to *P. aeruginosa* were performed at 25°C (Meisel, et al., 2014). In line with this, we observed more robust *daf-7p::gfp* expression in ASI neurons in P0 animals exposed to PA14 at 25°C (Figure 2). However, none of these adjustments resulted in robust elevated *daf-7p::gfp* expression in F2 progeny (Figure 2, Table 1, and Table S1). While 25°C growth enhanced PA14-induced *daf-7p::gfp* expression in P0 animals, the effect on the F2 generation was an apparent increase in variability, as the F2 progeny of PA14 and OP50 trained animals were more likely to show statistically significant differences in *daf-7p::gfp* expression, but in either direction (Figure 2). Similarly, modified training and growth conditions to enable inheritance of learned avoidance behavior did not result in significant changes to P0 and F1 inheritance results and did not result in robust detection of learned avoidance in the F2 progeny (Figure 3 and Table S2).

### Using a single-copy *daf-7p::gfp* reporter strain did not result in elevated *daf-7p::gfp* expression in ASI neurons among F2 progeny

The strain FK181 contains an integrated multi-copy *daf-7p::gfp* reporter and the co-injection marker *rol-6(su1000)* [pRF4] tandem array (Murakami et al., 2001). We noticed sporadic loss of both the Rol-6 phenotype and *gfp* expression from the line, suggesting either spontaneous transgene silencing or changes in the *daf-7p::gfp* reporter copy number. Similar instability has been noted in the Murphy lab (C. Murphy personal communication). To characterize this in greater detail, we established and maintained nine independent FK181 lines and observed co-incident loss of the Roller phenotype and *gfp* expression in all nine lines, and in six of nine lines this loss of array phenotypes was heritable (Table S3). It is known that growth conditions and mutations that modulate small RNA pathways can affect transgene expression levels from multi-copy tandem arrays (MacMorris et al., 1994; Cui et al., 2006; Fischer et al., 2013). Because this key reagent is not reliable, we obtained QL296, a strain containing an integrated single-copy *daf-7p::gfp* reporter that does not include a co-injection marker with a morphological or growth phenotype (Zhan et al., 2015). The QL296 fluorescent readout was less bright than FK181 but reproduced the elevated *daf-7p::gfp* fluorescence in P0 and F1 progeny. Encouragingly, the coefficient of variation was reduced to about 0.27 (Figure S4), which should increase the sensitivity of this assay to detect differences in *daf-7p::gfp* expression. However, measured *daf-7p::gfp* levels in F2 descendants of trained and control P0 animals carrying the single-copy reporter were usually indistinguishable, as we had seen with the multi-copy reporter (Figure 2).

### Performing the learned avoidance assay in light or dark conditions did not produce heritable F2 avoidance

A recent report indicates that visible light contributes to *C. elegans* ability to detect and avoid PA14 (Ghosh et al., 2021). We found that choice assays performed in the light (on benchtop) or in the dark (in a closed cabinet) both readily produced learned (P0) and inherited (F1) PA14 avoidance but that F2 inheritance of learned avoidance was not reliably detected in either assay condition (Figure 3 and Table 2). The choice index ([number of animals choosing OP50 - number of animals choosing PA14] / Total number of choices) for assays performed in the light for both trained and control animals were frequently higher than the choice index for assays performed in the dark, however the learning indexes (the relative differences) were indistinguishable. Thus, ambient lighting conditions can impact the measured choice index between control and PA14 trained animals, but they do not detectably disrupt the learning index. Overall, none of the environmental adjustments were sufficient to reliably reproduce the reported results; the summary analysis of the learning indexes for all experiments showed highly significant P0 training results, modest F1 intergenerational inheritance, and insignificant F2 transgenerational inheritance (Figure 3J).

### OP50 growth conditions strongly affect OP50 aversion

Naïve (OP50 grown) worms often show a bias towards PA14 in choice assays (Zhang et al., 2005; Ha et al., 2010; Moore et al., 2019; Pereira et al., 2020; Lalsiamthara and Aballay, 2022). This response, rather than representing an innate attraction to PA14, likely reflects the context of the worm’s recent growth on OP50, a mild *C. elegans* pathogen (Garigan et al., 2002; Garsin et al., 2003; Shi et al., 2006). Thus, the naïve worms presented with a choice between a recently experienced mild pathogen (OP50) and a novel food choice (PA14) initially choose the novel food instead of the known mild pathogen (OP50 aversion). Because the difference in the choice between trained and naïve animals in the P0 generation is highly significant, while the difference in the F1 generation is much reduced (Figure 3), even a slight reduction in naïve aversion to OP50 could affect the ability to detect PA14 avoidance in F1 and F2 animals. Indeed, in our experiments (Figure 3), the control worms frequently showed a choice index score higher than that reported by Moore et al., (2019) and Kaletsky et al., (2020). In line with our results, some other groups have also reported higher naïve choice index scores (Lee et al., 2017). This variability in naïve choice may reflect differences in growth conditions of either the OP50 or PA14 bacteria. In addition, we note that among the studies that show naïve worm attraction to *Pseudomonas* (OP50 aversion) there are extensive methodological differences from the methods in Moore et al., (2019; 2021b), including differences in bacterial growth temperature, incubation time, whether the bacteria is diluted or concentrated prior to placement on the choice plates, the concentration of peptone in the choice plates, the length of the choice assay, and the inclusion of sodium azide in the choice assays (Zhang et al., 2005; Ha et al., 2010; Moore et al., 2019; Pereira et al 2020; Lalsiamthara and Aballay, 2022). Thus, the cause of the variability across published reports is not clear. Furthermore, because OP50 pathogenicity is enhanced by increased *E. coli* nutritive conditions (Garsin et al., 2003, Shi et al., 2006), the growth of F1-F4 progeny on High Growth (HG) plates (Moore et al., 2019; 2021b), which contain 8X more peptone than NG plates and therefore support much higher OP50 growth levels, immediately prior to the F1-F4 choice assays may further contribute to OP50 aversion among the control animals. We note that in our hands, in each aversion assay experiment that produced a significant F1 result, the average F1 control choice index score was lower than that detected in the preceding P0 generation where the control animals were trained on NG plates (Figure 3). Thus, changes in growth conditions that enhance OP50 aversion (lower choice index score) could magnify the difference between trained and control animals.

To test the effect of OP50 growth conditions on OP50 aversion we plated young adult N2 animals from HG OP50 plates on either HG OP50 or NG OP50 plates prepared exactly as for control training plates. After 24 hours the HG OP50 “trained” and NG OP50 “control” animals were tested on standard OP50 vs PA14 choice assay plates. In four experiments the magnitude of the differences in mean choice index (Learning Index, LI) exceeded 0.4 (Figure 4). However, the inter-experiment variability was also high, with two experiments failing to detect a difference and two experiments showing an inverse result to the other four. We note that when analyzing the sum of all worm choices across all choice assay plates by Fisher’s exact test (Figure S5), five of eight experiments show that HG OP50 growth conditions induce OP50 avoidance, and seven of eight experiments show significant differences between worms cultured for 24 hours on NG OP50 and HG OP50 plates. Although the results in most experiments are consistent with the hypothesis that HG OP50 exposure enhances OP50 aversion (negative learning index), due to the variability between experiments we interpret the results as inconclusive. Even so, these results highlight the sensitivity of the choice assay to environmental differences and demonstrate the not inconsequential difference between NG OP50 and HG OP50 conditions. Although the experimental design implicitly controls for the difference between P0 and F1 generation growth conditions, the magnitude of the NG OP50 vs HG OP50 response, which can exceed the magnitude of the OP50 vs PA14 F1 response but in the opposite direction, suggests that this experimental variable may contribute to the irreproducibility of the published results. Indeed, any enhanced OP50 aversion that results from growing worms on HG OP50 plates likely increases the ability to detect lingering PA14 aversion.

**Figure 4.**
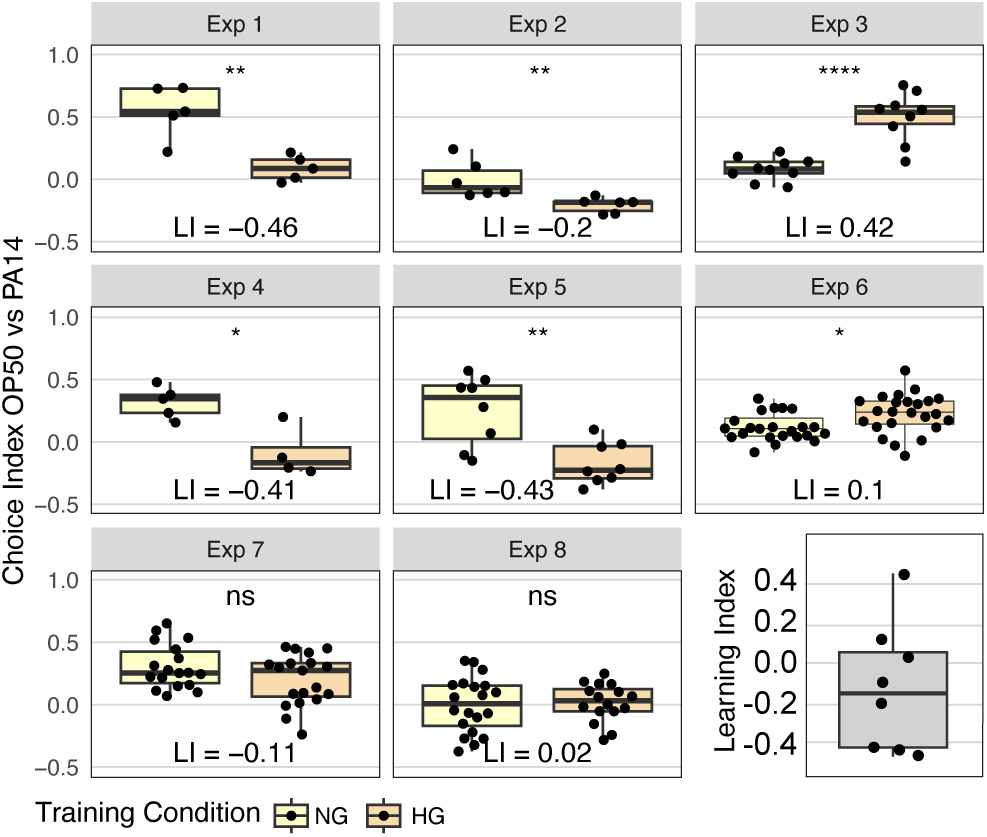
Effect of OP50 growth conditions on OP50 aversion. N2 worms grown to adulthood on HG plates were “trained” on either HG OP50 or NG OP50 plates for 24 hours and then assayed on OP50 vs PA14 choice plates. The choice index was calculated as described in Figure 1. The learning index, difference between mean HG choice index and mean NG choice index, for all eight experiments is plotted in the last panel. See Figure S5 for Fisher’s exact test analysis of choice assay results. Statical significance **** P < 0.0001, *** < 0.001, ** < 0.01, * < 0.05, ns > 0.05. See Methods section for statistical methods. Data for this figure is presented in Table S6.

While these observations do not explain our inability to replicate the published results, they do illustrate the sensitivity of the choice assay to differences in bacterial growth conditions. This supports the conjecture that non-obvious growth condition differences, beyond what we have explicitly tested, may contribute to the discrepancy between our results and the previously published results.

### The systemic RNAi pathway genes *sid-1* and *sid-2* act in parallel or downstream of the neuronal TGF-β pathway for intergenerational (F1) inheritance of pathogen avoidance

We were able to replicate the requirement, as reported (Kaletsky et al., 2020), for *sid-1* and *sid-2* in intergenerational (F1) inheritance of learned avoidance. Kaletsky et al., (2020) used a weak *sid-1(pk3321)* allele that remains fully sensitive to feeding RNAi targeting intestinal *act-5* (Whangbo et al., 2017), thus we repeated these experiments with either an early nonsense *sid-1(qt9)* allele (aversion assay) or a cas9 generated deletion of the entire coding region, *sid-1(qt158)* (*daf-7p::gfp* expression assay), both of which eliminate systemic RNAi silencing. We found that *sid-1(qt9)* P0 animals learn to avoid PA14 (learning index 0.58 and 0.43) but that their F1 progeny showed no inherited learned avoidance (learning index -0.04 and 0.00) (Figure 5A, B). Similarly, we found that *sid-2(qt42)* P0 animals also learn to avoid PA14 (learning index 0.87 and 0.67) while their F1 progeny showed little inherited learned avoidance (learning index 0.06, 0.08) (Figure 5C, D). Unexpectedly, the exogenous RNAi (*rde-1*), heritable RNAi (*hrde-1*), systemic RNAi (*sid-1*), and feeding RNAi (*sid-2*) pathways were not required for intergenerational inheritance of elevated *daf-7p::gfp* expression in the F1 progeny of PA14-trained animals (Figure 5E-H). Since Moore et al., (2019) showed that like *sid-1* and *sid-2* mutants, *daf-7* mutant P0 animals fail to transmit learned avoidance to their F1 progeny, we conclude that *sid-1* and *sid-2* must act in parallel or downstream of the neuronal TGF-β pathway for F1 inheritance of learned avoidance, not upstream as proposed by Kaletsky et al., (2020). If learned avoidance and the neuronal TGF-β pathway act in parallel, then the relative strength and time of activation of the two responses may be separately regulated. This is in line with our observation that identical training conditions (20°C) produce more reliable aversion behavior than *daf-7p::gfp* upregulation in the P0 generation yet more reliable *daf-7p::gfp* upregulation than aversion behavior in the F1 generation.

**Figure 5.**
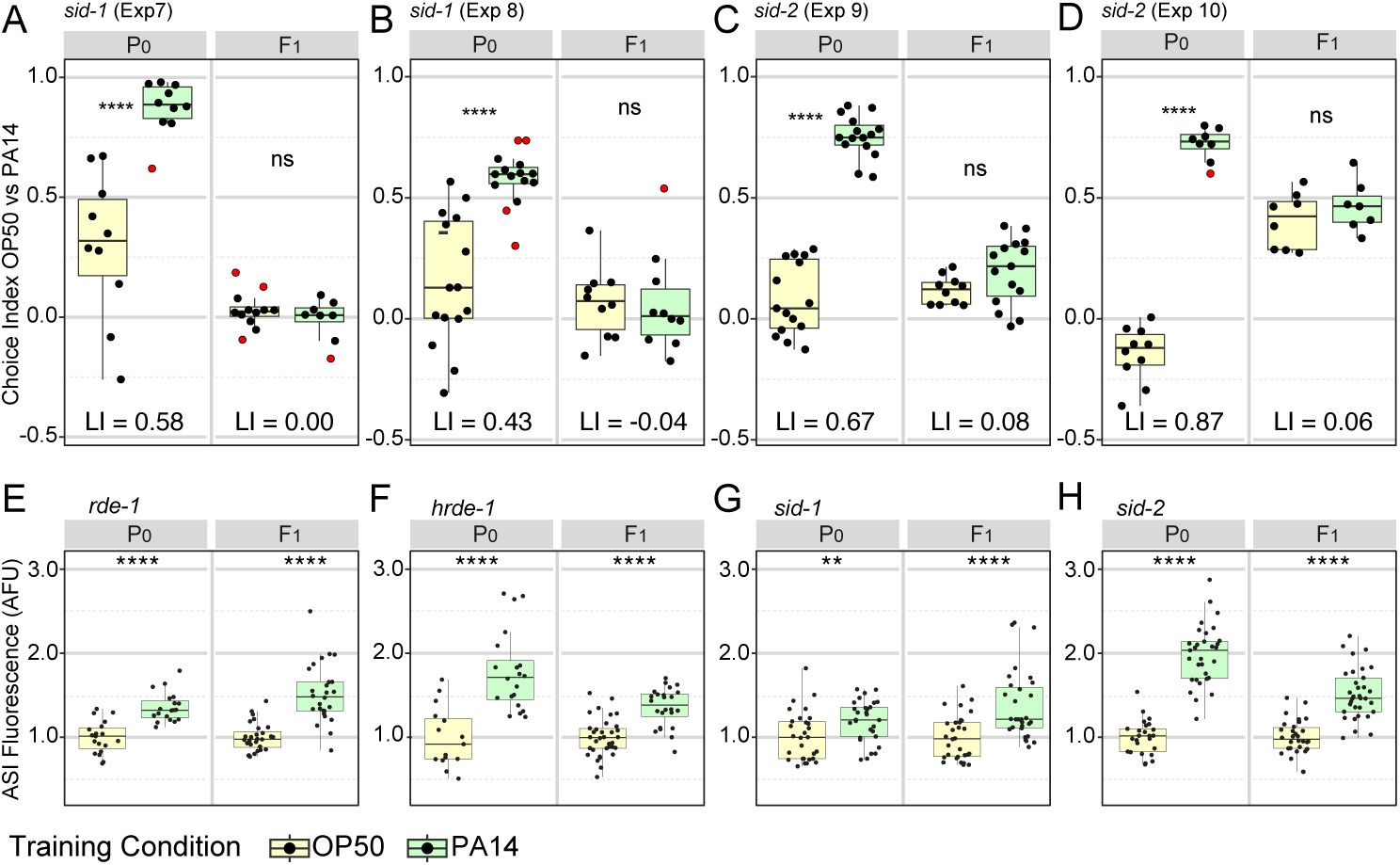
Effects of RNAi pathway mutants on intergenerational (F1) inheritance of avoidance behavior and ASI *daf-7p::gfp* expression levels. A-D, *sid-1(qt9)* and *sid-2(qt42)* choice assays. Growth and assay conditions for each experiment are described in Table 2. The choice index was calculated as described in Figure 1. See Figure S6 for Fisher’s exact test analysis of *sid-1* and *sid-2* choice assay results. N2 choice assay results performed in parallel with the *sid-1* and *sid-2* experiments are presented in Figure S7. E-H, ASI expression experiments showing average ASI neuron *daf-7p::gfp* expression levels per worm for each of the four genotypes are shown. ASI *daf-7p::gfp* expression levels are normalized to the average OP50 value by generation within each experiment for ease of presentation. The experiments shown in panels E-H were performed with animals cultured and trained at 20°C. Statical significance **** P < 0.0001, *** < 0.001, ** < 0.01, * < 0.05, ns > 0.05. See Methods section for statistical methods.

### Reanalysis of small RNA seq data identifies candidate siRNA-regulated pathogen response genes

The RNAi mutant analysis (Figure 5) confirmed the results by Moore et al., (2019) and Kaletsky et al., (2020) that *sid-1* and *sid-2*, which are required for systemic and feeding RNAi, are important for intergenerational inheritance of PA14 avoidance. Since antisense endo-siRNAs have been implicated in heritable gene silencing (Minkina and Hunter, 2018; Duempelmann et al., 2020), PA14 induced changes in small RNAs targeting mRNAs that respond to PA14 exposure may identify heritable RNAi-regulated pathogen response pathways. To investigate the contribution of regulatory small RNAs to heritable pathogen avoidance, Moore et al., (2019) sequenced mRNAs and small RNAs from control and PA14 trained P0 animals. The assembled small RNA sequencing libraries contain both sense-strand (piRNAs or 21U-RNAs and miRNAs) and antisense-strand (endogenous siRNAs) small RNAs, yet the published work appears to have presented only findings on the sense-strand small RNAs. To determine whether PA14-induced changes in antisense small RNA levels may contribute to heritable pathogen responses, we reanalyzed the sequencing data from Moore et al., (2019). We detected sense-strand small RNAs that correspond to 2041 genes that increased (1429) or decreased (612) by 2-fold or more (Padj. ≤ 0.05) (Figure 6A, Figure S8A), which, using our pipeline, is similar to the mapping data published by Moore et al., (2019) (1252 up and 450 down). We also mapped differentially expressed antisense small RNAs to over 4000 genes (3928 up ≥ 2-fold and 171 down > 2-fold (Padj. ≤ 0.05)) (Figure 6B, Figure S8B). We then plotted the predicted mRNA targets that changed by 2-fold or more (P adj. ≤ 0.05) against the antisense small RNAs that changed by 2-fold or more (P adj. ≤ 0.05), identifying 116 mRNAs that are putatively regulated by the endogenous RNAi pathway in response to PA14 exposure (Figure 6C). Curiously, the *maco-1* gene (red dot in Figure 6C), identified as the regulatory target for the piRNA mediated multi-generational response to PA14 exposure (Kaletsky et al., 2020), is not targeted by many differentially expressed antisense endo-siRNAs. Thus, the effect of the *P. aeruginosa*-expressed small RNA P11 on *maco-1* expression, which was identified in a subsequent study (Kaletsky et al., 2020), is not likely to be directly mediated by the endogenous RNAi pathway.

**Figure 6.**
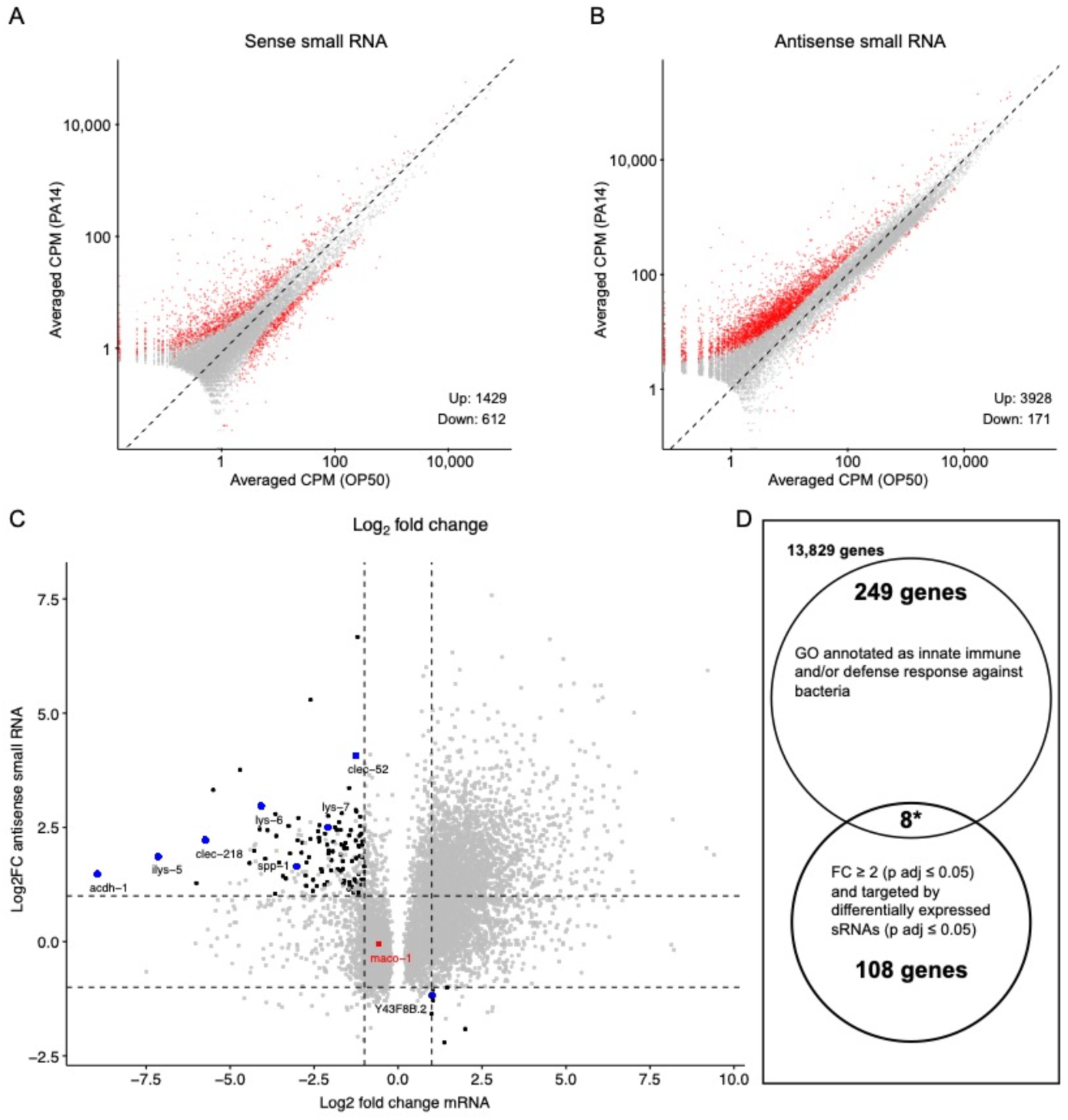
Re-analysis of small RNA and mRNA sequence data from PA14 exposed and control P0 animals. RNA sequence data (PRJNA509938) was downloaded and re-analyzed as described in Methods. A, B, Scatter plots comparing sense and antisense strand small RNAs (17-29 nucleotides) from worms grown on PA14 and OP50. Red dots represent ≥ 2-fold change and Benjamini-Hochberg corrected P values (P adj. ≤ 0.05) and grey dots correspond to less than a 2-fold change or insignificant difference (P adj. > 0.05). Volcano plots of the same data are shown in Figure S8. C, Black dots represent genes associated with small RNAs that show a significant 2-fold or greater change in abundance (Y axis) and map to an mRNA that also shows a significant 2-fold or greater change in abundance (X axis). Blue dots correspond to the subset of target mRNAs associated by GO analysis (panel D) with anti-bacterial or innate immune responses. The red dot represents the *maco-1* gene. D, Gene Ontology analysis of differentially expressed mRNAs targeted by differentially expressed small RNAs. Eight of the 116 small RNA targeted differentially expressed mRNAs are also associated with antibacterial or innate immune responses, while 257 of all 14,194 detectably expressed genes share similar GO annotations representing a 3.8-fold (P < 0.0001, hypergeometric distribution) enrichment over neutral expectations.

We then analyzed gene ontology descriptions of the detectably expressed genes (Schindelman et al., 2011) to determine whether any of the subset of genes potentially regulated by the endo-RNAi pathways are likely to contribute to heritable pathogen aversion. We found that 257 of the 14,194 detected expressed genes are annotated as either innate immune response or defense response against bacteria (Figure 6D). Eight of the 116 genes putatively regulated by endo-siRNAi pathways (blue dots in Figure 6C; 3.8-fold enrichment) are so annotated (Table S4). This analysis demonstrates that small RNA regulated genes are differentially expressed in response to pathogen exposure. While the identified genes are candidates for mediating a heritable pathogen aversion response, to support this conjecture it will be necessary to determine whether they and their candidate small RNA regulators are differentially regulated in the F1 and F2 progeny of pathogen exposed animals.

### Summary of thoughts and concerns regarding the potential causes of the irreproducibility

Likely causes for the discrepancy between our results and the published reports include uncontrolled environmental variables or genetic drift (microbes or worms). To control for this, we included in our investigation independent isolates of OP50 and PA14, we cultured worms at different growth temperatures, and we prepared bacterial samples under different growth regimes (aeration vs standing liquid cultures). We note that previous bacterial RNA sequence analysis identified a small non-coding RNA called P11 whose expression correlates with bacterial growth conditions that induce heritable avoidance (Kaletsky et al., 2020). Critically, *C. elegans* trained on a PA14 ΔP11 strain (which lacks this small RNA) still learn to avoid PA14, but their F1 and F2-F4 progeny fail to show an intergenerational or transgenerational response (Figure 3L in Kaletsky et al., 2020). The fact that we observed an intergenerational (F1) avoidance response is evidence that our PA14 growth conditions induce P11 expression. We also confirmed that our PA14 growth conditions induced strong pathogenicity (Figure S9).

Furthermore, we showed that OP50 control culture conditions can dramatically affect naïve worm behavior in the PA14-OP50 choice assay (Figure 4), supporting the conjecture that environmental factors may contribute to the observed discrepancy. Some environmental differences may be systemic and rooted to laboratory or geographical constraints, including humidity, which could affect salinity levels, lysogen activation, or the presence of other, potentially contaminating bacteria in the environment reflecting adjacent laboratory activities. If these potential environmental differences are sufficient to obscure the detection of the transgenerational effect, then the robustness and ecological significance of the transgenerational effect in a natural setting must be minor.

The imaging-based assay measures the expression of a multi-copy *daf-7p::gfp* transgene. We noted that although this transgene array is integrated, the FK181 strain must be monitored for maintenance of the Rol phenotype, which correlates with detection of *gfp*. Similar observations have been made by the Murphy group (C. Murphy personal communication). The instability of the FK181 strain may reflect sporadic transgene silencing or transgene-array rearrangements resulting in lower transgene expression. These observations illustrate that this key reagent is not reliable. To control for this, we obtained the published single-copy non-Rol *daf-7p::gfp* strain QL296 (Zhan et al., 2015). Using this single-copy *daf-7p::gfp* reporter we detected a robust difference in the P0 and F1 generations, but did not detect significant differences in the F2 generation (Figure 2, Table S1).

Ideally, we would have compared our results with the published results (Moore et al., 2019), to possibly identify additional experimental parameters for further investigation; for example, a quantitative comparison of naïve choice in the P0 and F1 generations could help to determine the role of bacterial growth in the choice assay response. However, none of the raw data for the published figures and unpublished replicate experiments (Moore et al., 2019) were available on the publisher’s website or provided upon request to the corresponding author. In the absence of a quantitative comparison, it remains possible that an explanation for the discrepancies between our results and those of Moore et al., (2019) has been overlooked.

## Concluding remarks

Although we cannot explain the cause of the differences between our results and the published results, we can confidently conclude that this example of TEI is, at present, insufficiently robust for experimental investigation of the mechanisms of multi-generational inheritance. This does not negate the possibility that future investigations may reveal a critical experimental or environmental variable that enables robust multigenerational inheritance. Our attempts to troubleshoot the protocol eliminated several variables, including growth temperature, developmental timing, genetic drift of either WT (N2) or reporter strains (FK181 and QL296) or *P. aeruginosa* strain PA14 (all refreshed from independent stocks). In addition, we determined that some environmental differences associated with the choice assay (light), although potentially altering the choice index, had similar relative effects on trained and control animals (the learning index). PA14 is a lethal pathogen and OP50 is a mild pathogen, thus the most likely cause for the discrepant results is a subtle difference in OP50 physiology in the post P0 generations that affects the magnitude of OP50 aversion. Indeed, we showed that 24-hour exposure to slightly more pathogenic OP50 can dramatically enhance OP50 aversion (Figure 4).

While we were unable to reliably replicate the reported F2 aversion or *daf-7p::gfp* expression responses, we confirmed that the F1 aversion response does require the dsRNA transporters *sid-1* and *sid-2*. While this remains a fascinating result, we would not presume that the transported substrate is therefore an RNA molecule, particularly because the systemic RNAi response supported by *sid-1* and *sid-2* is via long double-stranded RNA. To date, no evidence suggests that either protein efficiently transports small RNAs, particularly single-stranded RNAs (Feinberg and Hunter, 2003; Shih and Hunter 2011; McEwan et al., 2012). How these two proteins support intergenerational epigenetic inheritance remains a mystery. Unfortunately, since neither *sid-1* nor *sid-2* is required for the robust intergenerational increase in *daf-7p::gfp* expression in ASI neurons, the lack of a robust single-animal assay severely hampers mechanistic investigation using this paradigm.

The heritable behavioral and physiological plasticity induced by pathogens is likely to reveal fundamental evolutionarily and ecologically relevant pathways. However, the history of heritable epigenetic research has frequently been hobbled by the difficulty of controlling all environmental factors and the lack of reproducible phenotypic assays. Thus, independent reproducibility is of paramount concern, and we have tried to be completely transparent as a model for how heritability research should be presented within the *C. elegans* community.

## Methods

The protocol as described in (Moore et al., 2019; Kaletsky et al., 2020; Moore et al., 2021b; Sengupta et al., 2023) was followed closely, with a few clarifications and updates, some suggested in the Murphy lab updated protocol (C. Murphy pers comm), as detailed in the S1_aversion_protocol.

### Bacterial growth and plating

Bacterial cultures were seeded on NG (Normal Growth) or HG (High Growth) plates. NG media: per liter: 3 g NaCl, 17 g agar, and 2.5 g peptone in H_2_O, autoclave, cool to 55°C then add 1 mL cholesterol (5 mg/mL), 1 mL 1M CaCl_2_, 1 mL 1M MgSO_4_, 25 mL 1M KPO_4_ (pH 6.0) (Brenner 1974). HG media: per liter: 3 g NaCl, 30 g agar, and 20 g peptone in H_2_O, autoclave, cool to 55°C then add 1 mL cholesterol (5 mg/mL), 1 mL 1M CaCl_2_, 1 mL 1M MgSO_4_, 25 mL 1M KPO_4_ (pH 6.0) (based on Rose et al., (1982)). NG plates minimally support OP50 growth, resulting in a thin lawn that facilitates visualization of larvae and embryos. HG plates (8X more peptone) support much higher OP50 growth, resulting in a thick bacterial lawn that supports larger worm populations.

For the choice assay, OP50 for seeding HG growth plates prior to training was grown at room temperature (2 days) without aeration and used fresh or stored at 4°C for up to 1 month. Seeded HG plates were incubated at RT for 2-4 days before use or incubated for 2-4 days and stored at 4°C for up to 14 days. Only freshly grown OP50 (37°C with aeration, 16-22 hours) was used for aversion training and choice plates and was used at growth density or diluted in LB to OD 1.0. For the *daf-7p::gfp* ASI experiments, we used OP50 grown either at 37°C with aeration and diluted to 1.0 OD/ml or at room temperature without aeration for growth and training plates (see Table S1), again without effect on the results (Table S1, Figure 2, and Figure S2). PA14 for seeding training plates for both the avoidance assay and *daf-7p::gfp* ASI experiments and for preparation of choice assay plates was grown for 14-19 hours at 37°C with aeration and diluted to OD 0.5 or 1.0 in LB broth before plating. Whether PA14 cultures were grown for 14 or 18 hours had no discernable effect on pathogenicity (Figure S9). OP50 and PA14 seeded training plates and choice plates were incubated at 25°C for two days, and then equilibrated to room temperature before use. PA14 and OP50 training plates were incubated either in separate incubators and/or separate partially covered boxes within the same humidity controlled (<50% relative humidity) room during worm training.

### Worm growth and synchronization

Wild-type (N2, FK181, QL296) and mutant worms (Table S5) were grown on NG or HG OP50 plates without starvation, crowding, or contamination for a minimum of three generations before hypochlorite treatment to obtain P0 embryos. Fresh hypochlorite solution was prepared immediately prior to each use. Adult worms were pelleted or allowed to settle (15 mL tube), resuspended in 5-10 mL of hypochlorite solution, and, to minimize contact time with the hypochlorite solution, mixed continuously by nutating or vortexing until less than 5% of the adult body parts were visible. In some experiments, as the adults were initially breaking open (∼3-5 minutes) and the solution began to acquire a yellow tint, the worms and released embryos were pelleted, resuspended in fresh hypochlorite solution, and mixed for an additional 2-3 minutes, until less than 5% of adult body parts were visible. The released embryos were pelleted and washed 4X in 5-10 mL of M9 buffer. In the ASI assays, the last M9 wash optionally contained 0.01% Triton X100 to inhibit embryos from sticking to the plastic. We note that the published protocols assert that worms must not be centrifuged immediately prior to bleach treatment and that bleach-treated embryos must be handled gently. Our results do not support this assertion, as the adult worms bleached to prepare P0, F1, and F2 embryos for experiments 4.1 and 4.2 (Figure 3, Table S2) were treated gently or centrifuged and vortexed during bleaching respectively and showed no meaningful differences between the results.

In all experiments, animals from one or several pooled training plates were treated as a single biological replicate, with a portion of the animals assayed and a portion used for propagation for the next generation. We note that experiments 5.1 and 5.2, which were performed in parallel using a homogenous starting population that was split and then trained and assayed in parallel, showed statistical variation (Figure 3, Table 2).

### Measuring *daf-7p::gfp* levels in ASI neurons

Both wild-type *daf-7p::gfp* strains (FK181, QL296) and QL296-derived RNAi pathway mutants (HC1218, HC1221, HC1222, and HC1223) were maintained and trained on OP50 or PA14 as described above. Immediately before each respective imaging session approximately 30-40 trained or control animals were transferred by platinum wire directly to 5 µL of 2.5-10 mM levamisole in M9 on 10% agarose pads. A coverslip was added and sealed with beeswax. For the FK181 strain, GFP z-stacks were collected at 1 μm intervals at 63x magnification on a Zeiss spinning disc confocal microscope. For the QL296 strain, which has dimmer GFP than FK181, GFP z-stacks were collected at 0.5 μm intervals at 63x magnification on a Zeiss LSM 980 confocal microscope (Harvard Center for Biological Imaging). For data analysis, a maximum intensity projection (MIP) was generated for each z-stack and MIPs across all conditions and generations were then blinded. If the ASI neurons overlapped, then MIPs were generated from subsets of levels in the z-stacks without overlap and scored separately. Each ASI neuron was manually selected from each MIP using the ImageJ polygon selection tool (Schindelin et al., 2012). The mean pixel intensity was recorded from each and normalized to nearby background for each MIP. Although we also visually confirmed the expected PA14-induced *daf-7p::gfp* expression in the ASJ neurons of P0 animals, the fluorescence of these neurons was never quantified, as the ASJ response is not induced beyond the P0 generation (Moore et al., 2019). In the rare event that an ASI neuron was out of the frame or z-range in the collected z-stacks, the animal was excluded from analysis prior to quantification. We noted bimodality in the distributions of ASI *gfp* expression levels in our data, which likely reflects the position of the bilateral ASI neurons relative to the objective. To compensate for this, we used the average GFP level per pair of neurons in each worm (Figure 1, Figure 2, and Figure 5). We note that measured GFP expression levels in each ASI neuron within an animal are not independent, thus using the average avoids the artifactual bimodal distribution and reports the actual sample size for statistical purposes. We also note that coefficient of variation values using the average ASI level per worm were 20% lower for FK181 and 50% lower for QL296 (Figure S1) which should increase the ability to detect differences between trained and control animals. Although using the average for each neuron pair or treating each neuron as an independent sample altered the statistical significance of some experiments, it did not engender an F2 response (Figure S2). All collected data is presented in Table S1. Within each analysis pipeline, Z-scores were calculated from normalized intensity values for all samples within a data set, and any data point with a |Z| > 3 was removed. Unpaired, two-tailed Welch’s t-test was performed to generate all reported p-values. Statistics and figures were prepared using R Studio and refined using Adobe Illustrator (Wickham, 2016; Posit Team, 2023; R Core Team, 2023b; Wickham, 2023).

### Choice assay conditions, sample size, statistical analysis, and multigenerational propagation

Sodium azide is historically used to preserve transient worm choices in arena chemotaxis assays. In the food choice assay, the effect of the sodium azide can paralyze worms before they enter the bacteria lawn, possibly interfering with their choice. Azide also affects bacteria, potentially affecting the production of molecules that attract or repel worms and thus altering worm choice independent of the paralytic effect on worms. To determine whether sodium azide may influence worm choice, we performed some assays without sodium azide. We found that independent of the application of azide, most worms had made a choice within 30 minutes and that essentially no worm left their initial food choice during the first hour (data not shown). These no-azide assay plates were moved to 4°C after 30-60 minutes to preserve the initial food choice of the worms. After cold-induced rigor was achieved, individual assay plates were returned to room temperature for counting (S1_aversion_protocol). Individual worms were removed as counted to ensure complete and accurate counts. The addition of azide had no discernable effect on the choice assay results (Figure S10). Which method was used to preserve the worm’s initial choice is noted in Table 2, Table S2, Figure 1, and Figure 3.

For comparison to published results, we present the choice assay results in quartile box plots and report a Wilcoxon unpaired P-value for choice assays. We used a one-sample two-tailed T-test to calculate the P-value associated with the summary figure plotting the learning indices for each experiment (Figure 3J). The reporting of individual choice assay results as single data points in quartile box plots assumes that there are differences between choice plates that need to be included in significance tests and are independent of the number of worms assayed, limiting the power of the statistical analysis by the number of choice plates. Indeed, statements in Moore et al., (2019) and Kaletsky et al., (2020) suggest that some choice assay plates in these studies may have sampled as few as 10-20 worms per assay plate. However, under the assumption that there are no meaningful environmental differences between choice assay plates, the relevant outcome for the experimental design is the overall sum of choices across all choice plates. Here, the null model is that, independent of training condition, each worm chooses OP50 over PA14 with a probability preference p. Testing n worms, independent of the number of animals per choice plate or number of choice plates, the number that choose OP50 is X=pn, with X being binomially distributed. These results can be analyzed by a 2 X 2 contingency table: two conditions (OP50 vs PA14 training) and two outcomes (OP50 or PA14 choice). In this analysis, Fisher’s exact test can be used to test the effect of training on the probability preference p. The results of this alternative analysis of our data are presented in Figure S3, Figure S5, and Figure S6.

The Star Protocol (Moore et al., 2021b) cautioned that crowding may influence choice, but we found no difference between choice index scores and sample size (Figure S11). For each experiment (generation, training condition) we calculated Pearson’s correlation coefficient (r) for the number of worms per assay plate against choice index and then plotted r against the average number of worms per plate for each experiment.

R Studio was used for statistical tests and generation of graphical plots which were modified for presentation using Adobe Illustrator (Wickham 2016; Posit Team, 2023; R Core Team 2023; Wickham 2023).

### RNA-seq data analysis

PRJNA509938 RNA-seq data (Moore et al., 2019) were re-analyzed from the level of raw sequence reads. The reads were trimmed of universal adaptors using *cutadapt* (Martin, 2011) and aligned to the *C. elegans* WBcel235 genome assembly using *bowtie* aligner (Langmead et al., 2009) with no more than 2 mismatches allowed (v=2). Aligned reads were counted using *HTSeq-count* (Anders et al., 2015) (union mode) with settings *stranded=no* for mRNA-seq data and *stranded=yes* (for reads mapped to the same strand as the genomic feature) or *stranded=reverse* (for reads mapped to the opposite strand as the genomic feature) for small RNA data. Differential expression analysis was carried out using DESeq2 R package (Love et al., 2014). The absolute value of log2FC≥1 and Benjamini-Hochberg corrected p-value ≤0.05 were used to define significant changes in gene expression or small RNA levels. Volcano plots were built with EnhancedVolcano (Blighe, 2023) and ggpubr R packages (Kassambra, 2023).

### Pathogenicity assay

The Balskus and Murphy isolates of the PA14 strain were cultured at 37°C for 14 or 18 hours. OP50 was similarly cultured for 18 hours. Each culture was diluted in LB to OD 1.0 and 0.75 ml spread to cover the entire agar surface of NG plates which were then incubated at 25°C for 48 hours before addition of N2 adults. To prepare the N2 adults, HG OP50 grown worms (20°C) were bleached, and their embryos were placed on HG OP50 plates for 72 hours (20°C). A single population of adults were washed from the plates with M9, allowed to settle, washed once with M9, resuspended at 4-5 worms/µL and 20 µL spotted onto three technical replicate plates for each condition. The plates were then incubated at 20°C and scored at 24 hours and at 12-hour intervals thereafter until all the PA14 grown animals were dead. Corpses were counted and removed at each interval. Approximately 15-30% of the PA14 cultured worms were found desiccated on the walls of the plates at the 24-hour time point and were not included in the count. All living worms were transferred to freshly prepared plates at 48 and 96 hours. The fraction of surviving worms was determined by (1 – (cumulative number of dead worms at each scoring interval/sum of the number of living and cumulative dead worms at 120 hours)). All PA14 cultured animals were dead by 120 hours.

## Supporting information

S1_aversion_protocol

## Acknowledgments

We thank members of the Hunter lab for ongoing discussions and specifically Nicole Bush and Alexandra Weisman for detailed comments on the manuscript. We also thank L. Ryan Baugh for discussions and comments on the manuscript as well as Richard Losick and Sean Eddy for comments on the manuscript. This work was performed with the support of National Institutes of Health (NIH) grant GM089795 and through the support of Grant 62579 from the John Templeton Foundation. The opinions expressed in this publication are those of the authors and do not necessarily reflect the views of the John Templeton Foundation. Some strains were provided by the Caenorhabditis Genetics Center, which is funded by NIH Office of Research Infrastructure Programs (P40 OD010440). We thank the Harvard Center for Biological Imaging (RRID:SCR_018673) for infrastructure and support. We also thank WormBase.

## Author contributions

DPG Methodology, Formal Analysis, Investigation (*daf-7p::gfp* expression in ASI), Writing – Review and Editing

AVS Software, Formal Analysis (RNA-seq data reanalysis)

CPH Methodology, Formal Analysis, Investigation (aversion testing by choice assay), Writing – Original Draft, Review and Editing, Visualization, Supervision, Project Administration, Funding Acquisition.

## Supplemental information

S1_aversion_protocol

S2_supplemental_figures_tables: Figures S1 – S11, Tables S3 – S5

Data Tables S1, S2, S6, S7, S8. Excel files containing data related to Figures 1 – 5, S1 – S11.

