## Supplementary material for "Reported transgenerational responses to *Pseudomonas aeruginosa* in *C. elegans* are not robust": S1_aversion_protocol

### PA14 training and choice assay protocol

This is Craig Hunter's distillation and clarification of protocols to train and assess learned and inherited PA14 avoidance in N2 animals through the $F_2$ generation. Green highlights indicate known methodological difference from the Star Protocol (Moore et al., 2021).

**Materials**

Non-starved[1] N2[2] maintenance plates (20°C), Method A or B.

Method A, every three days transfer four L4 hermaphrodites to a new small (60 mM, vented) NG plate seeded with 0.15 – 0.30 mL of maintenance OP50[3]. Seeded and dried (2-4 days bench top) plates can be stored (tightly covered) at 4°C for 4 weeks. Warm to room temperature before use.

Method B, every ~2 days chunk a small (5-7 mm) portion of agar with worms/embryos to a new NG small OP50 (maintenance) seeded plate.

Up to 30 Large (100 mm) HG plates for growing $P_{-1}$ through $F_2$ populations.
8 Large NG plates for training.
90 Small (60 mm) NG plates for choice assays.
2 Large LB plates for streaking -80°C PA14 and OP50 glycerol stocks to single-colonies. You will require multiple single colonies to start fresh cultures over a ~10-day period.
5% Sodium hypochlorite (NaClO) (J.T. Baker) stored in the dark at 4°C (less than 3 months old; replace earlier if the resulting embryo prep displays low viability).
5 N KOH (stored in plastic, strong hydroxide will etch glass).
1 M $NaN_3$ (sodium azide), store small (50µL) aliquots at -20°C.
M9 buffer (3 g $KH_2PO_4$, 6 g $Na_2HPO_4$, 5 g NaCl, 1 mL 1 M $MgSO_4$, $H_2O$ to 1 liter. Sterilize by autoclaving).

The timing of this 20-day protocol is organized around **Day 0**, the day $P_0$ embryos are bleach synchronized. The timing of steps before **Day 0** does not need to be as precise as the steps after **Day 0**.

**Day -9**

[Seed $P_{-2}$ HG plates]

- Seed three to four large HG plates with 1 mL of maintenance OP50[4] spread over the entire surface. Leave to dry/grow on bench until Day -7.

**Day -7**

[Prepare $P_{-2}$ adult plates]

- Chunk a quarter of a mature (non-starved, mixed staged) maintenance NG small plate each to two to three freshly seeded large HG plates. Grow worms at 20°C[5].
  .

---

[1] A minimum of 3 consecutive generations without starvation; in practice, often greater than 10.
[2] The lab N2 stock (CGC reference 257) is refreshed from frozen stocks 2-3 times per year.
[3] Worm maintenance OP50: either a single colony from a fresh (<3 weeks) LB plate or a scraping from a -80°C glycerol stock is added to 400 mL of LB in a 500 mL bottle, incubated (without shaking) at 37°C overnight and then stored at 4°C for 4-8 weeks.
[4] Altering how the OP50 is prepared for pre-$P_0$ HG plates has not altered our results.
[5] 15°C growth prior to the $P_0$ generation (as described in Moore et al., (2019) and Kaletsky et al., (2020)) does not alter the behavior of trained or control animals.

- Monitor daily and discard contaminated plates, excessively crowded plates (worms crawling on sides or lids or showing dauer behavior - towers, tail standing), starved plates, and plates with more than 1/200 males.

**Day -6**

[Prepare P-1 HG plates]

- Seed three to four large HG plates with 1 mL of maintenance OP50 spread over the entire surface. Leave to dry/grow on bench until Day -3.

**Day -4**

[Streak fresh PA14 and OP50 from glycerol stock]

- Streak OP50 and PA14 to single colonies on their own LB plates; incubate 15-20 hours at 37°C.

**Day -3**

[Store PA14 and OP50 colony plates]

- Seal colony plates with parafilm and store inverted at 4°C (Colony plates should be re-streaked from glycerol stock if more than 2 weeks old).

[Obtain P-1 embryos by bleaching]

- Using M9 buffer, wash and pool adult worms from HG plate(s) into a 15 mL plastic centrifuge tube, let settle until most adults are in bottom ~1 mL of tube (settling helps remove younger larvae still suspended in supernatant). Aspirate/replace M9 solution once and after second settle reduce volume to 1 ml or less.
- Add 5-10 mL of freshly prepared[6] (0-2 hours) alkaline hypochlorite (bleach) solution (6.7 mL of M9 (or $H_2O$), 2.3 mL of 4-6% NaClO, 1.0 mL 5 N KOH)[7]. Nutate (rock gently) for 4-5 minutes. At about 5 minutes use a dissecting microscope to observe worms; they should start bending and rupturing at the vulva. Optionally mix solution and continue nutating[8].
- After this initial 4-5 minutes observe nutating worms every 30-60 seconds under dissecting scope. When only embryos and a few body parts (pharynx) are visible (usually 7-8 minutes), pellet embryos (30 seconds, #3 1400 rpm [~250g] IEC clinical centrifuge), aspirate bleach solution, add 10-15 mL M9[9] (wash 1). Pellet the embryos, aspirate the solution, and add fresh

---

[6] The Moore et al., 2021 procedures note that the prepared solution may be stored at either room temperature or at 4°C for up to a month. Hypochlorite solutions (stock and alkaline diluted) are sensitive to light and warm temperature, and prolonged storage may reduce the reagent's potency. Our stock 4-6% NaClO solution is stored in the dark at 4°C and replaced quarterly. To ensure consistency between experiments, we used only freshly prepared bleach solution.

[7] We note differences between the bleach recipe that we use and those used in Moore et al., (2019) and Moore et al., (2021). The differences between the bleach recipes in Moore et al., (2019) and Moore et al., (2021) indicate that variation in bleaching procedures is unlikely to impact the response to PA14. All recipes use a 4-6% stock solution of NaClO (assumed 5% to simplify comparisons). Our recipe produces an alkaline hypochlorite solution (1.15% NaClO, 0.5 M KOH) that when prepared and used fresh produces embryos ready for washing after 7-8 minutes. The recipe in Moore et al., (2019) produces a hypochlorite solution that is 1.5% NaClO and 0.25 M KOH, while the recipe in Moore et al., (2021) produces a hypochlorite solution that is 0.6% NaClO and 0.25 M KOH. Moore et al., (2021) notes using a 5 N KOH pH 6.0 stock solution, which is likely a misunderstanding as the pH of 5 N KOH is ~13.7. Hydroxide is added to diluted hypochlorite to increase the pH, stabilizing the NaClO for storage; addition of a low pH reagent would increase the degradation rate of the solution.

[8] Vigorous mixing can accelerate the bleaching process and reduce total contact time with the bleach solution. Vigorous mixing of embryos does not affect their behavior as adults (Figure 3, Gainey et al.).

[9] The increase from three to four washes and increase in wash volume from 2 mL to 10 mL more efficiently removes bleach solution, maximizing embryo recovery. Bleach smell should not be detectable in the last wash.

M9 three more times (4 washes total). Count the number of embryos in 2-3 samples (20-50 µL) of the third wash. Remove M9 from the fourth wash leaving enough buffer for ~20 embryos/µL.
- Plate 50 µL (800-1200 embryos) to each OP50 seeded HG plate (minimum of 3 plates). Drop the embryos into 3-5 distinct locations on the plate midway between the center and edge. Tilt and rotate the plate to let the solution of embryos spread over a larger surface area to accelerate the absorption of M9 and dilute any remaining bleach solution. Once absorbed, invert HG plates and place in an uncovered box in a 20°C incubator.
- Monitor as described above for $P_{-2}$ adults (**Day -7**).

[Prepare OP50 for $P_0$ HG plates]
- [17:00 hours (5PM)] Pick a single OP50 colony into 4 mL of LB, aerate in rotator for 16 hours at 37°C.

**Day -2**

[Prepare $P_0$ HG plates]
- [09:00 hours] Measure OD600 of 1:10 dilution of the OP50 culture, dilute the ON culture in LB to OD = 1.0, seed 1 mL to each of six to eight large HG plates. Tilt and rotate the plate to spread the OP50 over the entire surface.

**Day 0**

[Obtain $P_0$ embryos by bleaching]
- [23:00 hours (11PM)] Bleach the $P_{-1}$ adults to obtain $P_0$ embryos. Bleach, wash, count, and plate 2800 embryos per plate on six to eight HG plates as described above for $P_{-1}$ embryos (**Day -3**).
- The bleached synchronized $P_0$ embryos must develop for 56-60 hours prior to PA14 training[10]. The time of training will determine the time of the $P_0$, $F_1$, and $F_2$ choice assays for the next week (72-hour intervals). My preference is to bleach the $P_0$ eggs at 23:00 (Monday is convenient to avoid choice assays on a weekend) which allows an 8-9AM start time (Thursday) for training and choice assays (Friday, Monday, Thursday).

[Prepare PA14 and OP50 overnight (ON) cultures for training plates]
- [17:00 hours] For OP50, pick a single colony into 4 mL of LB. For PA14, pick two single colonies (insurance against biofilm formation) into two tubes with 4 mL of LB. Place tubes in rotator for 16 hours in 37°C incubator. Put eight large NG plates on bench (to be seeded the next day).

**Day 1**

[Seed training plates with OP50 and PA14]
- [09:00 hours] Briefly vortex the OP50 culture and one of the PA14 cultures without obvious biofilm[11], measure OD600 of 1:10 diluted sample. Add appropriate amount of ON culture to LB (5 mL) to make OD = 1.0 culture.
- Add 1 mL of each OD = 1.0 culture to each of four large NG plates. Rotate and tilt the plates to distribute the culture to cover the entire surface.
- Place lid up in 25°C incubator in humidity controlled[12] BSL2 room. Any condensation will form on the lid rather than cause excessive plate wetness.

---

[10] PA14 training slows development and production of $F_1$ embryos. The additional ~8 hours of growth beyond the recommendation in the Star Protocol enables consistent recovery of sufficient $F_1$ progeny and does not noticeably alter P0 learned avoidance (Figure 1 and Figure S1, Gainey et al., 2024).

[11] I have noticed a biofilm (stringy clumps) in such a PA14 culture only once in over 200 cultures.

[12] We employ a humidity-controlled environment (< 50%) to minimize seasonal environmental variation. The published protocol (Moore et al., 2021) mentions that wet plates can interfere with PA14 learning and choice.

[Prepare PA14 and OP50 ON cultures for choice assay plates]

- [17:00 hours] Pick two OP50 single colonies into two tubes of 4 mL of LB and two PA14 single colonies into two tubes of 4 mL of LB. Place tubes in rotator for 16 hours in 37°C incubator.
- Put 30 small NG plates on a bench. Mark the position for OP50 and PA14 spots on the bottom of each plate (small dots, different colors, approximately 5mm from the edge and 5 mm above the midline) and place an X at the worm origin position (this will form an equilateral triangle between the origin and the two food spots). These plates will be seeded on Day 2.

**Day 2**

[Seed P0 choice assay plates]

- [09:00 hours] Briefly vortex the two OP50 cultures and one of the two PA14 cultures without obvious biofilm, measure OD600 of 1:10 diluted sample. For choice plates, add appropriate amount of ON culture to LB (1mL) to make OD = 1.0 culture.
- Spot 25 µL of either OD = 1.0 OP50 or PA14 culture on appropriate spot on all 30 plates. I use a manual pipettor (not a repeat pipettor) to more gently deliver each 25 µl drop, which reduces splatter and irregular spot shapes. When the first spot is dry (~15-30 minutes), spot the other culture. When the second spot is dry, move the plates, with their lids up, to a 25°C incubator in the humidity controlled BSL2 room.

[Seed HG plates for F1 embryos]

- From the second OP50 culture prepare 7-9 mL OD=1.0 and spread 1 mL on each of six to eight large HG plates as described above.

**Day 3**

[Transfer P0 animals to training plates]

- [09:00 hours] 56-60 hours after plating of P0 embryos, transfer training plates from 25°C to bench (~21°C). After 10-15 minutes verify with infrared thermometer that plates are equilibrated to room temperature.
- Check P0 plates for contamination and verify that most worms are young adults (YA). Gently add 12 mL of M9 to the first P0 plate, swirl gently to minimize bacteria in the solution as you dislodge worms from the food. Gently pour M9 and worms to second plate, swirl, pour to third plate, swirl, and pour into 15 mL polypropylene tube (can pour over a plate lid to collect any drops). Pouring is much quicker and less stressful for the worms than transferring with a pipette.
- Let worms settle to bottom 1 mL of M9 and then aspirate most of the M9. Add M9 to 10 mL, remove and count worms in three small aliquots while the remaining worms are settling. Aspirate most of the M9. Repeat the wash if necessary to minimize OP50 carryover. The M9 solution should be clear and the worm pellet readily visible.
- Add M9 to adjust to 80 worms per µL[13] (based on average of three counts above). Using a wide-bore tip (or a tip trimmed with a razor blade) plate 10 µL (800 worms) on to each of four OP50 training plates and 40 µL (3200 worms) to each of four PA14 training plates. Tilt and rotate plates to hasten drying of solution and distribution of worms on the plates.
- Place training plates, lid up, in 20°C humidity controlled BSL2 room.

---

[13] Counting the worms ensures that enough worms are plated for subsequent steps and avoids crowding/starvation.

**Day 4**

[Perform and score $P_0$ choice assay]

- [08:30 hours] Transfer choice plates (25°C, 48 hours) to bench. Prior to beginning the assay, examine each plate and discard if contaminated or if either choice spot is irregular in shape, too close to the edge, or if splatter colonies are present. Shuffle the remaining plates to randomize the position of the plates in incubating box relative to the trained worm populations to be placed on them. Mark the wall of the plates (I use a stripe from different colored sharpie) to indicate whether OP50 or PA14 trained animals will be placed on them.
- Remove lids from PA14 choice plates and arrange for ease of spotting azide[14].
- Thaw sodium azide aliquot.
- Prepare fresh bleach solution and add 7 mL to two 15 mL tubes.
- Add 12 mL of M9 to each of two labeled (PA14, OP50) 15 mL tubes.
- Check conditions on each training plate to verify uniform growth, presence of laid embryos, and absence of contamination. Set aside any compromised plate.

**Process PA14 trained animals onto choice assay plates and $F_1$ growth plates**

- [09:00] Gently pour 12 mL of M9 onto the first PA14 training plate, swirl and pour to the second, and to each additional plate. Pour contents of last plate (combined contents of all PA14 training plates[15]) back into labeled 15 mL tube and let worms settle (~2-4 minutes).
- Aspirate, add 10 mL of M9 solution, let settle, repeat until M9 solution is clear (usually 1X-2X, but can take 3 washes if more training plates are used or if the swirling to dislodge worms removed excessive amounts of bacteria).
- While worms are settling add 1 µL of azide to center of each choice spot on choice plates to be used with PA14 trained worms. Replace lids.
- When transferring worms use a wide-bore tip or a razor blade to trim pipette tip (2-3 mm) to increase the size of the opening[16].
- Remove M9 from the last wash, leaving about 50 µL of solution above each 100 µL of worms. Flick tube to distribute worms in solution, pipette 5-7 µL of washed worms onto origin (X) of first choice plate, check on dissecting scope to estimate number of worms (~150-250[17]). Adjust volume and repeat for all plates, flicking the tube of worms before each withdrawal[18]. This should be performed quickly (6-10 seconds per plate). Note the start time.
- Add bleach solution to the remaining PA14 trained worms, nutate, wash, count embryos as described for **Day -3**.
- Add ~1000 PA14 trained $F_1$ embryos to each of 3-4 large OP50 seeded HG plates (from day 2) (labeled PA14, $F_1$ embryos). Place in 20°C incubator for 72 hours.

**Repeat as above for OP50 trained animals.** (Can be done in either order)

---

[14] These plates will only be used for the choice assay, so there is no concern about contamination for the ~15 minutes the lids are off.

[15] The combining of all trained worms to a single population eliminates any plate-to-plate variability. The wash and bleach procedures are streamlined to minimize the time between washing worms from training plates to the addition of bleach. If performed without distraction or interruption the trained animals will be suspended in M9 before addition of bleach for less than 8 minutes (including all washes). If multiple pools are to be analyzed in parallel, then process each pool individually.

[16] This will reduce shear forces on worms being spotted onto each choice plate.

[17] The accuracy of population-based assays is improved by increasing the numbers. We have not detected an effect on choice from using between 100 – ~600 worms per choice plate (Figure S11, Gainey et al., 2024).

[18] I place each choice plate on the dissecting scope stage, add the worms, check the number of worms, replace the lid, and load the next plate.

**Score choice assay plates**
After 60 minutes, count the worms[19] at each spot within 2-worm-lengths of the food edge (the azide will paralyze many worms before they enter the food, but this does not appear to significantly affect the measurement of choice) (Moore et al., 2021). If worms are paralyzed more than 2-worm-lengths from the food spots or if worms are moving within the food spots – discard the plate. Count “censored” worms not at origin (damaged/sick) or at either spot. Typically, less than 1% of all worms will be censored. If greater than 10% then exclude the plate from the experiment. If healthy worms remain at the origin, discard the plate.

As an alternative to using azide as a paralytic, choice plates can be moved to 4°C after 30-60 minutes to preserve the initial food choice[20]. The plates should be moved to a cold surface, agar side down, to quickly bring about rigor. If your refrigerator only has wire racks or plastic surfaces, introduce a glass or metal plate as a heat-sink surface. If the worms are still mobile after 5 minutes, check that the refrigerator is cold enough. You can also use a cold room. Once rigor sets in, the worms will remain immobile for hours. However, the rigor is fully reversible on return to room temperature; thus, to count choice plates, remove individual plates from 4°C and quickly count all the worms. Worms will begin moving after 3-4 minutes but leaving food is rare during the short counting interval[21]. Because the worms are moving, it is essential to remove worms as counted rather than marking their position with a marker. When done as described here, we detected no discernable difference in choice index between the two methods (Figure S10, Gainey et al.).

[Prepare PA14 and OP50 ON cultures for F1 choice assay plates]
- As described above (**Day 1**).

**Day 5**
[Seed F1 choice assay plates]
- As described above for P0 choice plates (**Day 2**).

[Seed six to eight HG plates for F2 embryos]
- As described above for F1 embryos (**Day 2**).

**Day 7**
[Perform F1 choice assay]
- As described above for P0 choice assay (**Day 4**), except worms are growing on HG plates.

[Prepare PA14 and OP50 ON cultures for F2 choice assay plates]
- As described above (**Day 1**), except only one OP50 culture is required.

**Day 8**
[Seed F2 choice assay plates]
- As described above for P0 choice plates (**Day 2**).

---

[19] I use a P200 pipette tip attached to an aspirator to remove worms as they are counted; this avoids double-counting worms. Change tips between plates or more frequently if it becomes clogged with bacteria. Can use a pick to separate closely packed worms prior to aspirating.
[20] This was originally done as a control to determine whether azide could be affecting the choice index.
[21] Count and remove worms on the periphery of the food spot first. We have not observed animals leaving the food before counting is completed. If reanimated worms begin leaving either food spot before counting is completed, discard the plate.

**Day 10**

[Perform $F_2$ choice assay]

- As described above for $P_0$ choice assay (**Day 4**), except worms are growing on HG plates and the $F_2$ adults do not need to be bleached and propagated unless $F_3$ animals will be assayed (in which case you should proceed as described for $F_1$ animals [Days 4-8]).
