## Supplementary material for "Reported transgenerational responses to *Pseudomonas aeruginosa* in *C. elegans* are not robust": S2_supplementary_data

**Supplementary data Gainey et al.**

**Table S3. FK181 genetic instability**

| | | Rol Progeny | | non-Rol Progeny | |
|---|---|---|---|---|---|
| FK181 line | Starve-chunk cycle with first observed non-Rol | GFP+ | GFP- | GFP+ | GFP- |
| 1 | 4* | | | | |
| 2 | 4 | | | | +++ |
| 3 | 7 | + | | + | +++ |
| 4 | 7 | +++ | | | |
| 5 | 7 | +++ | | | |
| 6 | 7 | + | | (+)† | +++ |
| 7 | 7 | + | | + | |
| 8 | 8 | 241 progeny \| 22% non-Rol, 81% GFP- | | | |
| 9 | 8 | 167 progeny \| 26% non-Rol, 46% GFP- | | | |

FK181 was grown to starvation (6 days, 20°C) on a small NG plate and then chunked to a fresh small NG plate and grown to starvation. The first observed non-Rol hermaphrodite (day 3) was a picked to a fresh plate the Rol and GFP phenotypes of the progeny recorded.
* Sterile
† weak GFP

**Table S4. Putative endo-siRNA targeted genes implicated in host-pathogen responses.**

| Gene | Summary description of predicted and/or documented activities |
|---|---|
| Y43F8B.2 | Predicted to enable kinase regulator activity and protein kinase A binding activity. Involved in innate immune response. |
| ilys-5 | Predicted to enable lysozyme activity. Predicted to be involved in defense response to Gram-positive bacterium. |
| acdh-1 | Predicted to enable acyl-CoA dehydrogenase activity. Involved in defense response to Gram-negative bacterium and innate immune response. |
| clec-218 | Involved in defense response to Gram-positive bacterium. |
| lys-6 | Predicted to enable lysozyme activity. Predicted to be involved in innate immune response and signal transduction. |
| lys-7 | Involved in defense response to other organisms. |
| clec-52 | Predicted to enable signaling receptor activity. Involved in defense response to Gram-positive bacterium. |
| spp-1 | Enables pore-forming activity. Involved in defense response to other organism and pore formation in membrane of another organism. Part of pore complex. |

Summarized descriptions from WormBase (WS291).

**Table S5.** Worm strains used

| Strain name | Genotype | Reference |
|---|---|---|
| N2 | WT | Brenner, 1974 |
| FK181 | *ksIs2 [daf-7p::gfp + rol-6(su1006)]* | Murakami et al., 2001 |
| QL296 | *drcSi89 [daf-7p::GFP; unc-119(+)]* | Zhan et al., 2015 |
| HC445 | *sid-1(qt9)* | Winston et al., 2002 |
| HC306 | *sid-2(qt42)* | Winston et al., 2007 |
| HC1221 | *rde-1(ne219); drcSi89* | This study |
| HC1222 | *hrde-1(tm1200); drcSi89* | This study |
| HC1223 | *sid-1(qt158); drcSi89* | This study |
| HC1218 | *sid-2(qt42); drcSi89* | This study |

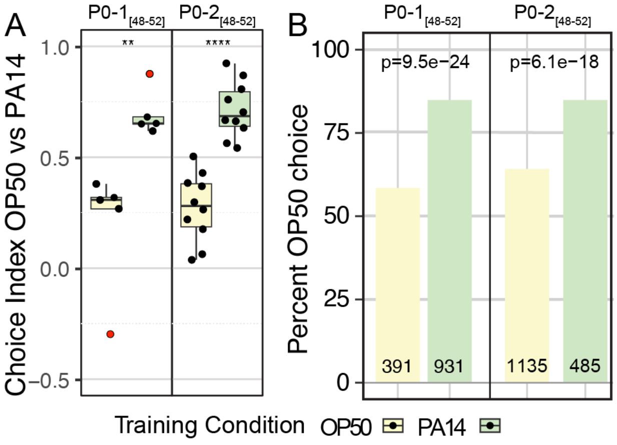


**Figure S1. P0 generation response to PA14 exposure.**
Results from two independent experiments in which training of $P_0$ animals began 48-52 hours post bleach and too few $F_1$ embryos were recovered to continue the experiment. A, Quartile box plots and training condition showing distribution of choice index values for each individual choice assay plate. Red dots indicate outlier data points that were included in all statistical tests. B, The same data as presented in panel A but expressed as a proportion of all individual choices across all assay plates. For both PA14 training and OP50 control animals the total number of OP50 and PA14 choices were summed across all choice plates and analyzed using a 2X2 contingency table and Fisher's exact test. Plotted is the percent OP50 choice for each condition (the PA14 choice is 100% minus the OP50 choice). The number of OP50 choice and PA14 choice animals scored for each experiment and training condition is indicated on each bar plot. The p-values represent the probability that the two observations came from identical, binomially distributed populations. These two experiments were performed with 20°C grown worms, the Balskus lab PA14 isolate was used, $P_0$-1 was performed in the light, $P_0$-2 in the dark and in both cases cold-induced rigor was used to preserve their choice. Statistical significance **** $P < 0.0001$, ** $P < 0.1$. See Methods section for additional statistical methods. Data for this figure is presented in Table S2.

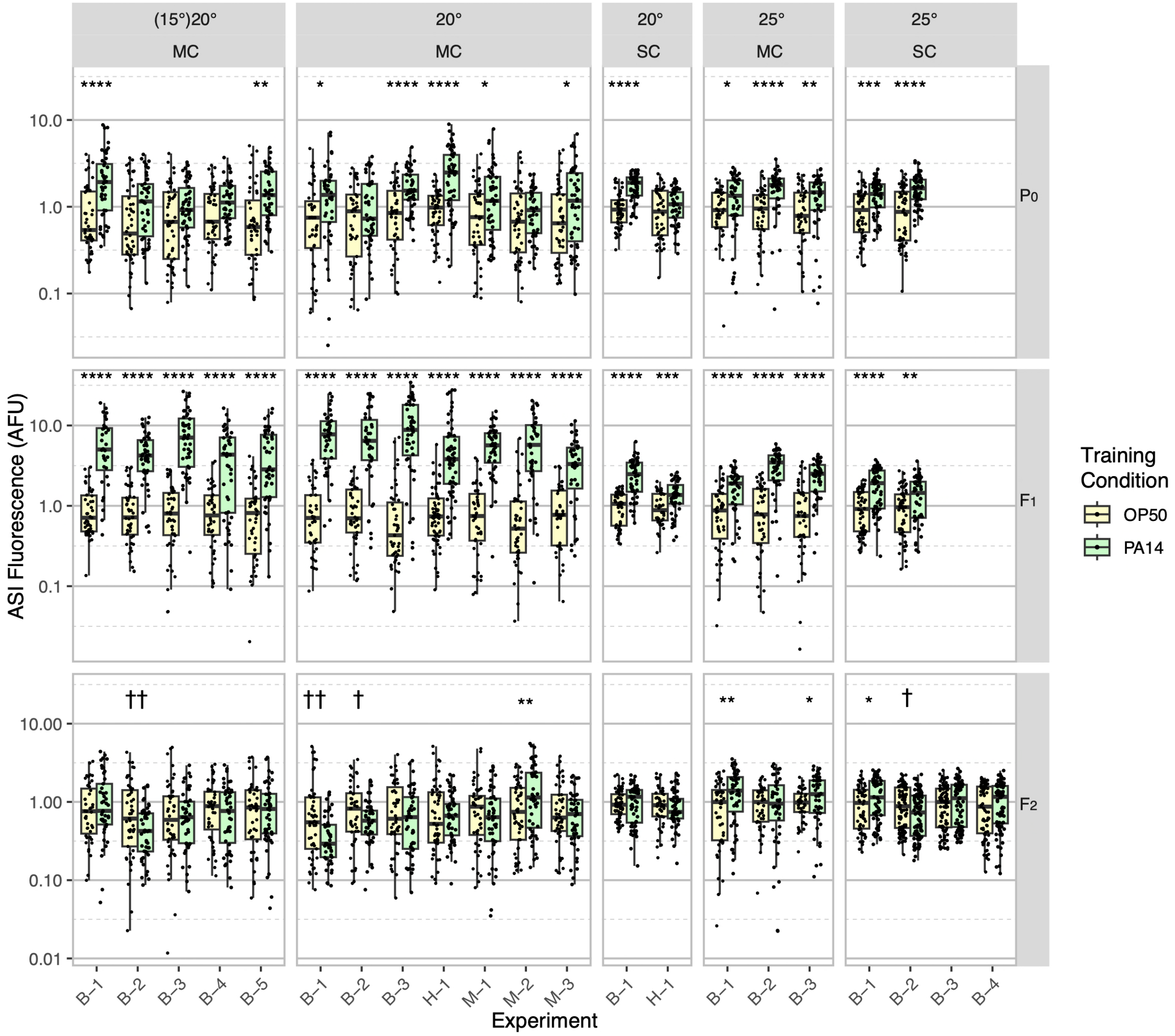


**Figure S2. Box plot display of individual ASI *daf-7p::gfp* expression levels after P0 PA14 exposure.**
Results from 21 independent experiments are normalized to the average OP50 value by generation within each experiment for summary presentation (experiments are named by experimental conditions, see Table 1). This figure includes the exact same data as shown in Figure 2 but plotted for all individual neurons rather than the average of the neuron pair in each animal. MC refers to FK181, which contains an integrated multicopy tandem array composed of the *daf-7p::gfp* reporter and the co-injection marker *rol-6(su1006)*. SC refers to QL296, which is a single copy insert of *daf-7p::gfp* with *unc-119(+)* as the co-selection marker (Zhan et al., 2015). Worms were cultured at either 20°C or 25°C and exposed to one of three different PA14 isolates (B, Balskus; H, Hunter; M, Murphy labs). In some experiments worms were grown at 15°C for at least three generations prior to the P0 generation (indicated with parentheses, i.e. (15)20). Statical significance **** P < 0.0001, *** < 0.001, ** < 0.01, * < 0.05. Non-significant labels (p > 0.05) are omitted for clarity. † Indicate statistical significance with control higher than experimental. See Methods section for statistical methods.

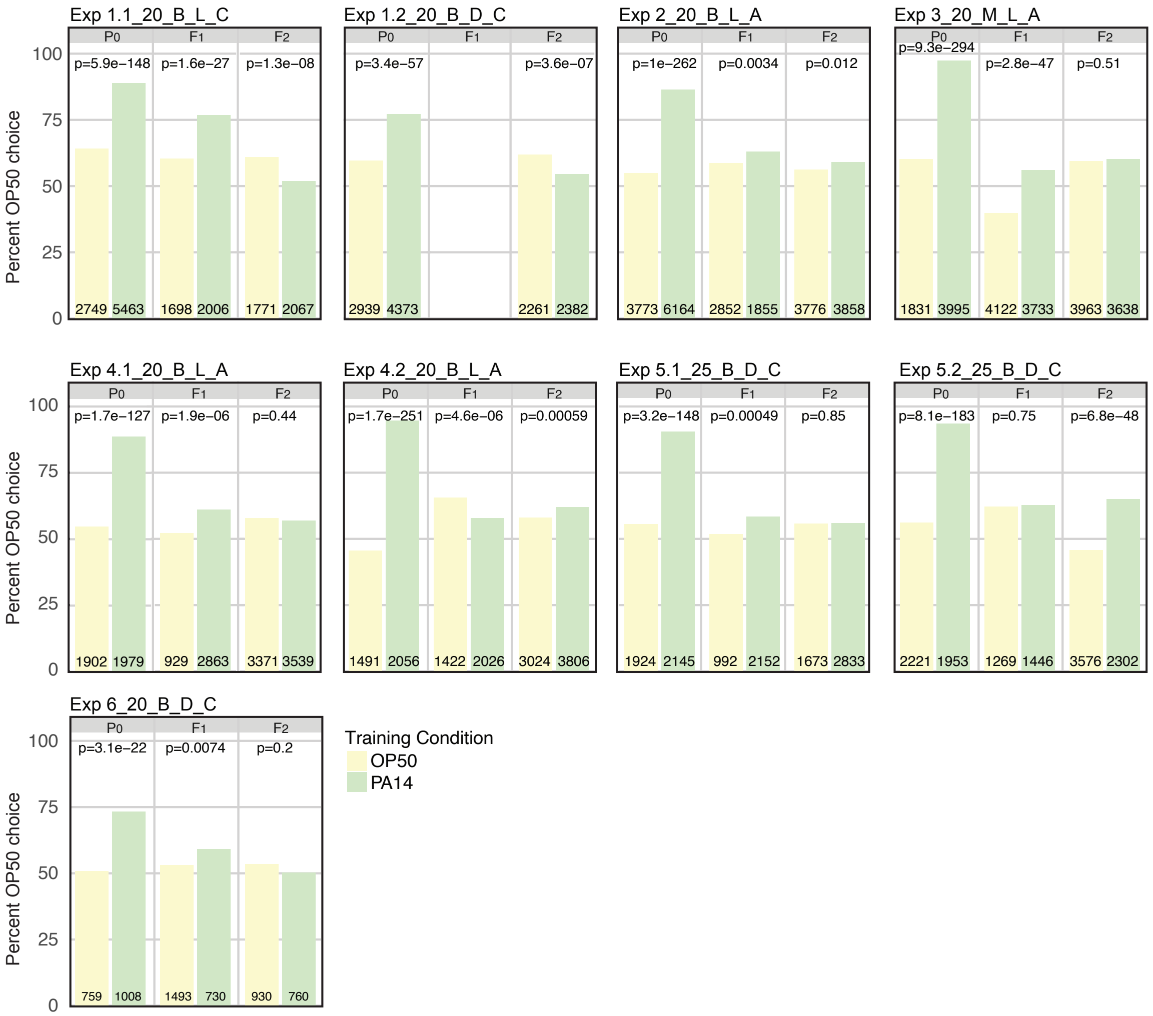


**Figure S3. P0 PA14 exposure fails to reproducibly induce F2 PA14 avoidance behavior.** The same data as presented in Figure 3 choice index plots. The experimental and assay conditions are indicated in the panel titles; [Exp #] [growth temperature (20°C or 25°C)] [PA14 isolate (B or M)] [light/dark assay condition (L or D)] [azide/cold paralytic (A or C)]. At each generation and for both PA14 training and OP50 control animals the total number of OP50 and PA14 choices were summed across all choice plates and analyzed using a 2X2 contingency table and Fisher's exact test. Plotted is the percent OP50 choice for each condition at each generation (the PA14 choice is 100% minus the OP50 choice). The number of OP50 choice and PA14 choice animals scored for each experiment, generation, and training condition is indicated on each bar plot. The p-values represent the probability that the two observations came from identical, binomially distributed populations.

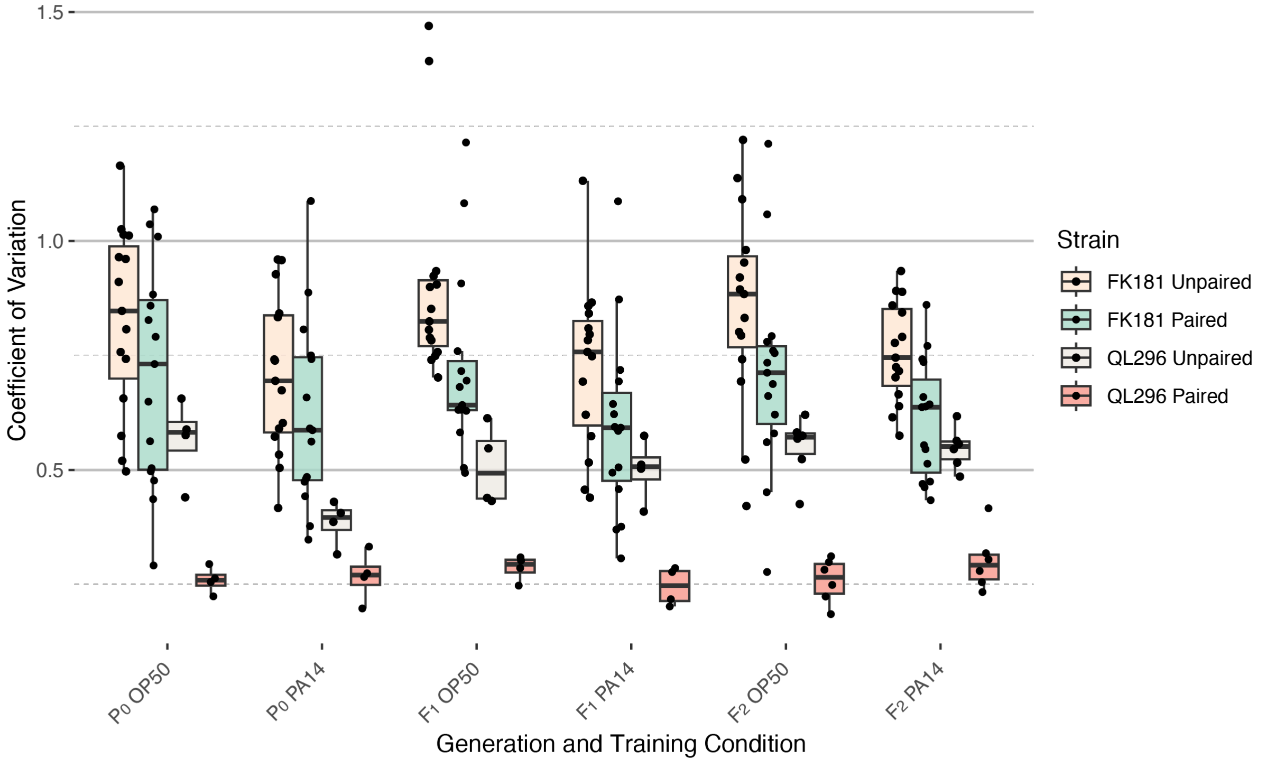


**Figure S4. Coefficient of variation analysis of ASI *daf-7p::gfp* expression levels.** The coefficient of variation was calculated for all experiments shown in Figure 2 (average of both ASI neurons - Paired) and Figure S2 (individual neurons - Unpaired) for both FK181 (integrated multicopy array) and QL296 (integrated single copy insertion).

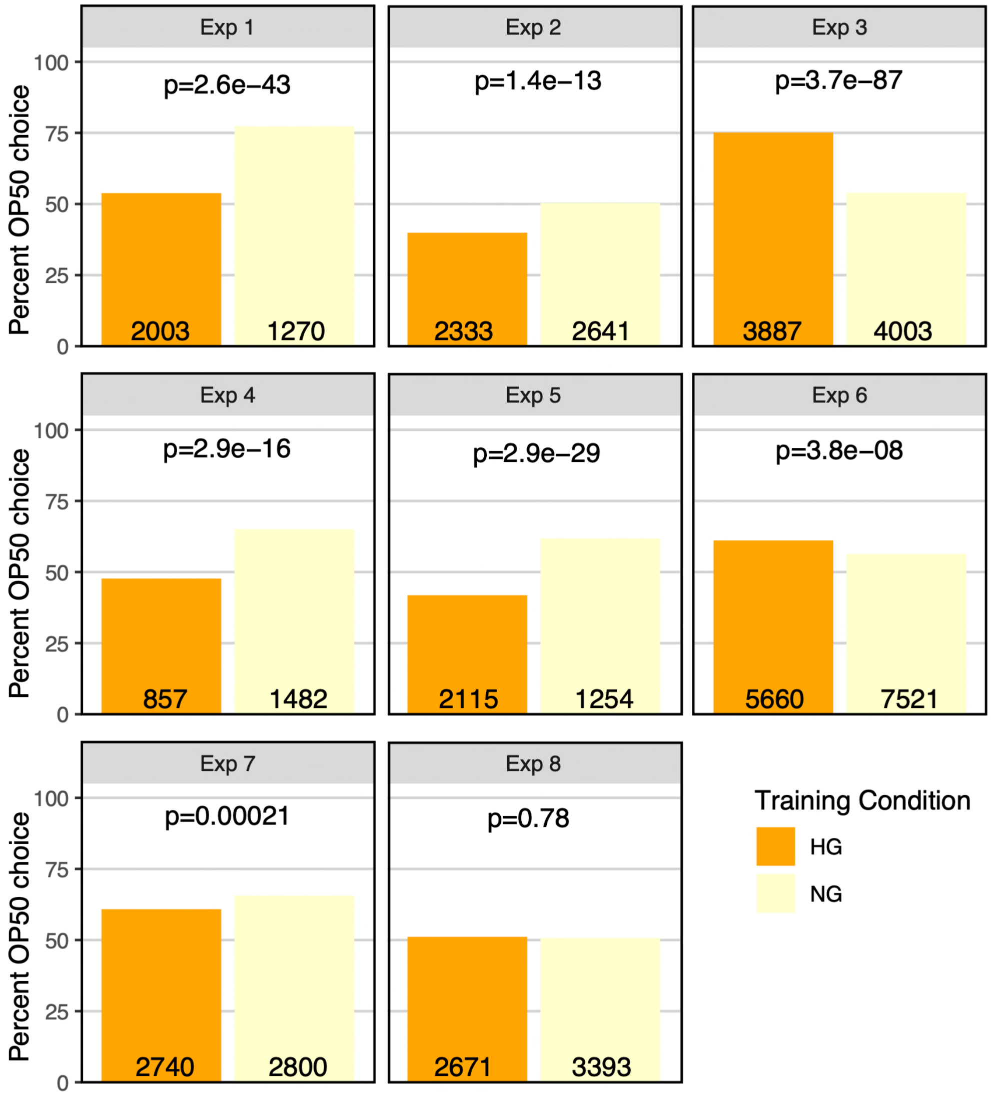


**Figure S5. OP50 growth conditions, independent of PA14 exposure, can affect OP50 vs PA14 choice.**
The same data as presented in Figure 4 choice index plots. N2 worms grown to adulthood on HG plates were "trained" on either HG OP50 or NG OP50 plates for 24 hours and then assayed on OP50 vs PA14 choice plates. For each of eight experiments the total number of OP50 and PA14 choices were summed across all choice plates and analyzed using a 2X2 contingency table and Fisher's exact test. Plotted is the percent OP50 choice for each condition (the PA14 choice is 100% minus the OP50 choice). The number of OP50 choice and PA14 choice animals scored for each experiment and training condition is indicated on each bar plot. The p-values represent the probability that the two observations came from identical, binomially distributed populations. Data for this figure is presented in Table S6.

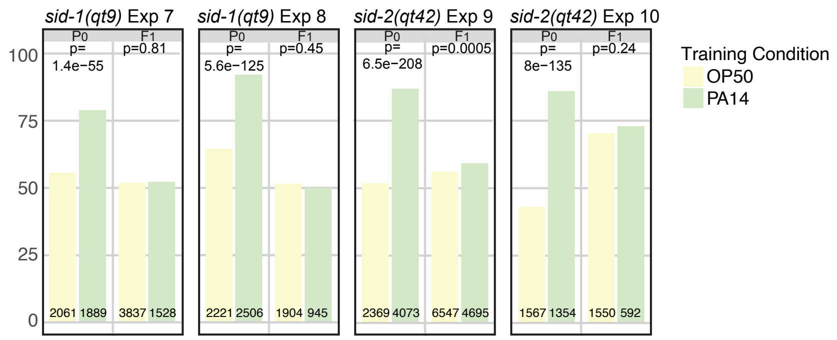


**Figure S6. *sid-1* and *sid-2* are required for intergenerational (F1) inheritance of avoidance behavior.**
The same data as presented in Figure 5 choice index plots. At each generation and for both PA14 training and OP50 control animals the total number of OP50 and PA14 choices were summed across all choice plates and analyzed using a 2X2 contingency table and Fisher's exact test. Plotted is the percent OP50 choice for each condition at each generation (the PA14 choice is 100% minus the OP50 choice). The number of OP50 choice and PA14 choice animals scored for each experiment, generation, and training condition is indicated on each bar plot. The p-values represent the probability that the two observations came from identical, binomially distributed populations.

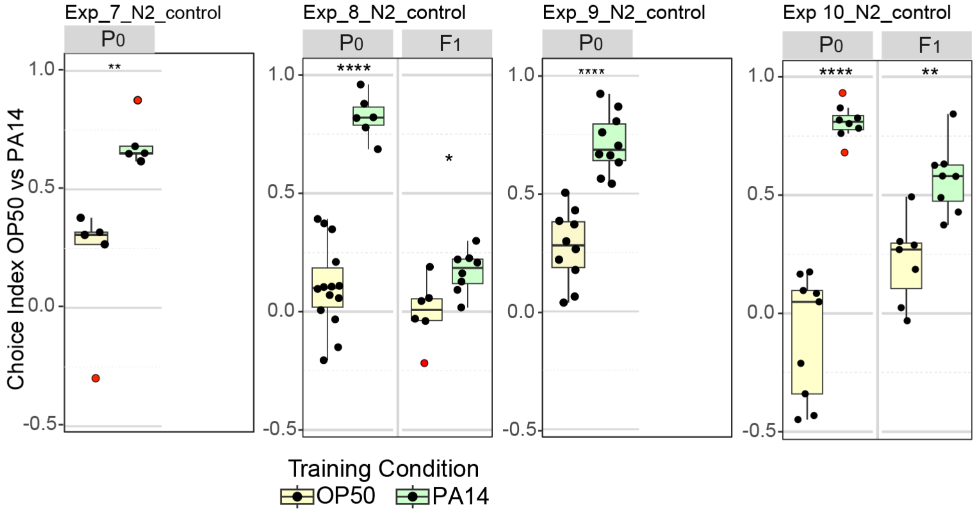


**Figure S7. N2 $P_0$ and $F_1$ choice assay results performed in parallel to *sid-1* and *sid-2* experiments (Figure 5).**

Choice assay results for N2 animals grown, trained, and assayed in parallel to experiments 7-10 (Figure 5). Experimental conditions are as described for experiments 7-10 (Table 2). Choice index calculated as described in Figure 1. Too few F1 animals were recovered in N2 controls for experiments 7 and 9 to perform the choice assays. Data for Exp_7_N2_control and Exp_9_N2_control are also shown in Figure S1. Data for Exp_8_N2_control is also shown in Figure 3, Exp 5.1. Statistical significance **** $P < 0.0001$, *** $< 0.001$, ** $< 0.01$, * $< 0.05$, ns $> 0.05$. See Methods section for statistical methods. Data for this figure is presented in Table S2.

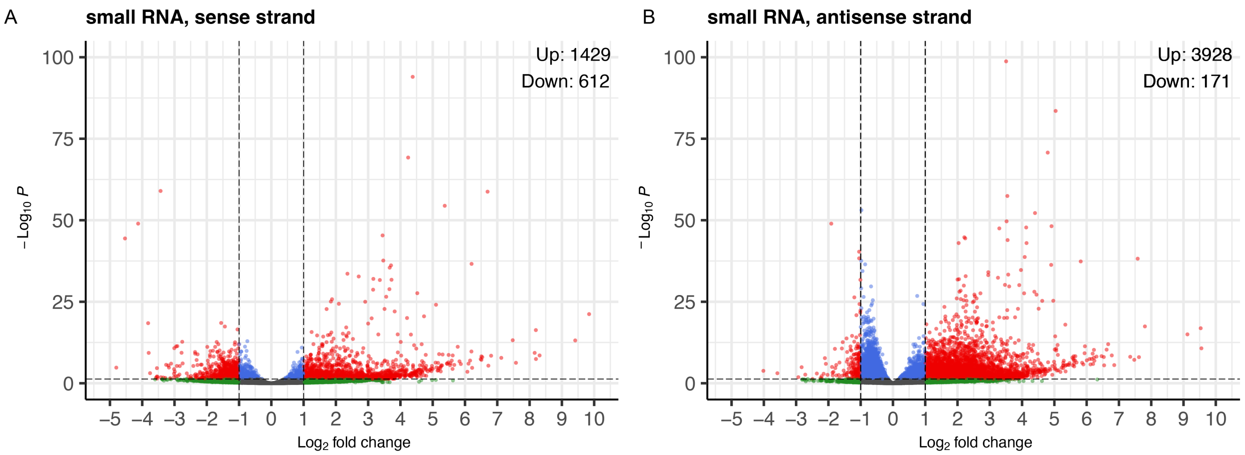


**Figure S8. Volcano plot displays reanalyzed small RNA sequence data from PA14 exposed and control P0 animals (Figure 6).**

RNA sequence data (PRJNA509938) was downloaded and re-analyzed as described in methods. A, B, PA14/OP50 Log(2) fold change for sense and anti-sense strand small RNAs (17-29 nucleotides), color coded for fold change and Benjamini-Hochberg corrected P values (P adj.). Small RNAs were mapped to genes for fold-change and P-value calculations. Black dots correspond to less than a 2-fold change and insignificant difference (P adj. > 0.05). Blue dots correspond to significant differences (P adj. ≤ 0.05) but less than 2-fold change. Green dots correspond to greater than 2-fold change and insignificant difference (P adj. > 0.05). Red dots represent small RNA clusters that show a 2-fold or greater significant difference (P adj. ≤ 0.05). Volcano plots were generated with the EnhancedVolcano R package (Blighe et al., 2023).

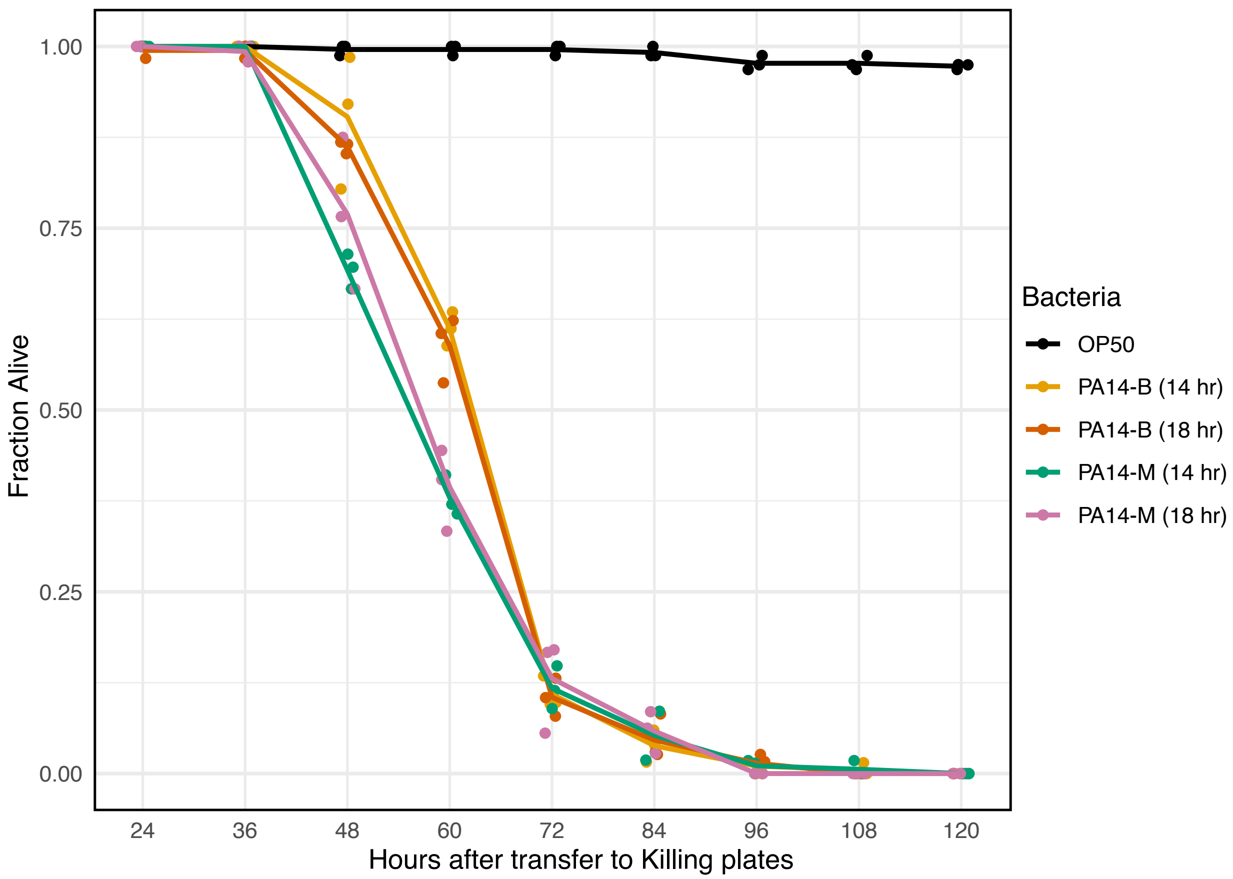


**Figure S9. The pathogenicity of PA14 isolates.**
Day 1 N2 adults were transferred to plates seeded with the indicated 14-hour or 18-hour bacterial cultures and then scored for viability at 12-hour intervals. PA14-B and PA14-M refer to the Balskus and Murphy lab PA14 isolates, respectively. Data points were jittered along the X-axis for display. n= 38 – 80 per replicate, 3 replicates per condition. Data for this figure is presented in Table S7.

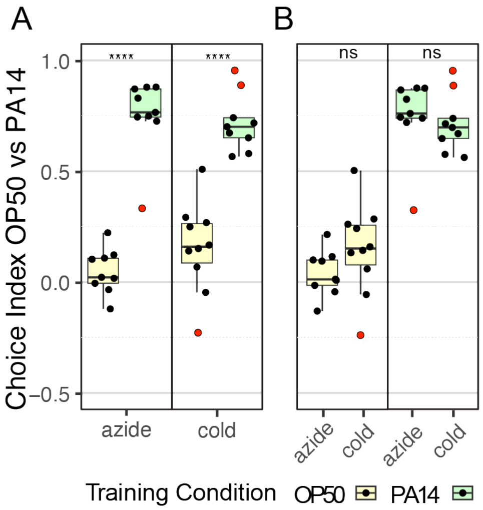


**Figure S10. Comparison of paralytic treatments.**
Quartile box plots showing distribution of choice index values for each individual choice assay plate for the indicated paralytic treatment and training condition. The $P_0$ generation of a single biological replicate (Exp 2, Figure 1, Table 2, and Table S2) was assayed on choice assay plates; half the choice plates were treated with azide and half were treated with cold-induced rigor. Red dots indicate outlier data points that were included in all statistical tests. Panel A compares the control and training response and statistics for each paralytic. Panel B compares the paralytic response and statistics for each training condition. Statistical significance **** P < 0.0001, ns > 0.05. See Methods section for statistical methods.

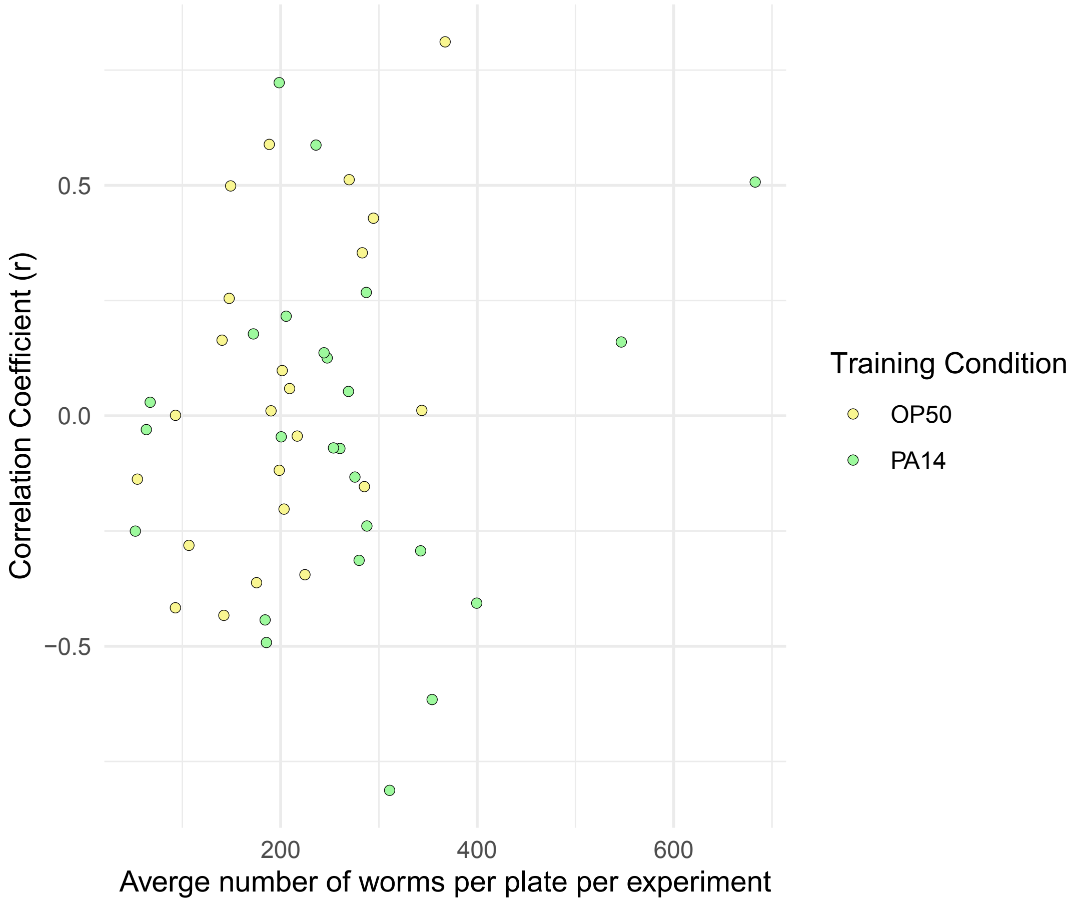


**Figure S11. Sample size does not correlate with choice index scores.**
The correlation coefficient (r) calculated from sample size vs choice index for each experiment (generation, training condition) shown in Figure 3 is plotted against the average number of worms in each experiment. Experiments with six or fewer choice plates were excluded. Data for this figure is presented in Table S8.
